# Adaptive brain rewiring after brain injury via adult neurogenesis

**DOI:** 10.64898/2026.09.22.753668

**Authors:** Oisorjo Chakraborty, Samuele Maturi, Michael Zabolocki, Mary Muhia, Sabrina Villar-Pazos, Meghane Decroocq, Simone Horenkamp, María Nazareth González Alvarado, Lukasz Piszczek, Matthias Schlachter, Tereza Oppelt Duranova, Henrik Skibbe, Jürgen A. Knoblich, Sofia Grade

**Affiliations:** Institute of Molecular Biotechnology of the Austrian Academy of Sciences (IMBA), Vienna BioCenter (VBC), Vienna, Austria; Vienna BioCenter PhD Program, Doctoral School of the University of Vienna and Medical University of Vienna, Vienna, Austria; Institute of Science and Technology Austria, Klosterneuburg, Austria; Institute for International Strategy and Emerging Technologies, Tokyo International University, Japan; Brain Image Analysis Unit, RIKEN Center for Brain Science, Wako, Saitama, Japan; Department of Informatics, Faculty of Informatics, Matsuyama University, Matsuyama, Ehime, Japan

## Abstract

The adult mammalian brain has limited regenerative capacity yet retains substantial potential for functional reorganization after experience, injury or disease^1–3^. While local plasticity at injury sites has been described^4,5^, the brain-wide consequences of a focal injury and the biological processes that drive them remain unknown. To address this, we performed an unbiased, whole-brain screen of neuronal activity at single-cell resolution following a focal injury to the primary visual cortex (V1) in mice. Olfactory processing areas emerged as loci of remote activation. We further show that V1 injury promotes the recruitment of adult-born neurons into the olfactory bulb (OB) and potentiates cortical feedback onto bulbar circuits. Combining high-density multielectrode array recordings with two-photon calcium imaging revealed OB circuit refinement characterized by increased synchrony and sharpened tuning of principal output neurons. These circuit-level functional enhancements coincide with improved behavioral performance in odor-guided tasks. Motor cortex lesions do not elicit similar cellular and behavioral responses, suggesting that such distal adaptation only develops when injury is coupled with increased olfactory demand. Together, these data demonstrate that the adult brain can recruit alternative sensory circuits distant from a focal lesion to undergo adaptive, functionally relevant reorganization, and implicate adult neurogenesis as a contributing mechanism. Our findings expose a previously underappreciated degree of remote plasticity and reveal a novel role of adult neurogenesis in sensory compensation.

## Main text

Clinical and experimental observations suggest that neural systems can compensate for substantial perturbations through mechanisms collectively referred to as neural plasticity. In neurodegenerative disorders, such as Parkinson’s and Alzheimer’s diseases, symptoms emerge only after extensive neuronal loss or neuropathology, indicating that compensatory processes can preserve function for prolonged periods^6,7^. Sensory deprivation can trigger functional reassignment of cortical territories and enhanced processing in spared modalities^8^. These examples illustrate the resilience of neural circuits in the central nervous system and their ability to reorganize in response to challenges.

Studies on brain injury have revealed that neural plasticity extends beyond changes in the strength of existing synapses. Neurons can undergo axonal sprouting^5^, form new synaptic connections^9^, and newly generated neurons are recruited to injured circuits^10–12^. Such observations have led to the view that the adult brain retains a latent capacity for structural remodeling. However, studies to date have focused on local responses within or near the injured tissue, leaving the true magnitude of induced plasticity at the whole-brain level unresolved. Fundamental questions remain: what are the brain-wide consequences of a focal lesion, which biological mechanisms drive remote circuit reorganization, if any, and do such changes contribute to functional adaptation?

Here, we generated a systems-level understanding of how neural networks respond beyond the site of injury. We combined unbiased whole-brain activity mapping at single-cell resolution with circuit, cellular, and behavioral analyses to investigate the consequences of a focal injury to the primary visual cortex (V1) in mice. This approach enabled us to identify remote regions recruited following injury and determine the mechanisms and functional significance of their reorganization.

### Remote activation of olfactory circuits after visual cortex injury

To determine the brain-wide consequences of a focal cortical injury, we evaluated changes in neuronal activity patterns as an indicator of circuit plasticity. We performed a stab wound injury (SW) in the V1 of adult mice and quantified neuronal activity across the entire brain 8-10 weeks later (Fig. 1a). Activity-dependent labeling was achieved using the targeted recombination of active populations (TRAP) method^13^, which relies on a *Fos*-CreER^T^^2^ knock-in mice crossed to a Cre-dependent reporter line. Administration of 30 mg/kg 4-hydroxytamoxifen (4-OHT) enabled permanent tdTomato labeling of cells active in the defined post-injury time point (Fig. 1a) compared to uninjured controls, in behaving animals in their home cage. tdTomato-positive cells were registered to a mouse brain reference atlas to generate brain-wide activity maps. This unbiased analysis revealed widespread changes in brain activity following V1 injury (Fig. 1b, left). As expected, visual areas showed reduced TRAP cell counts (Fig. 1c), consistent with persistent effects of the local injury on V1 integrity and recruitment. Interestingly, several multisensory integration areas ranked among the regions showing the largest increases in TRAP labeling, with cell counts ranging from 3-to 7-fold higher than those in intact mice (Fig. 1b, right). This finding motivated us to further investigate sensory systems across the brain. Analysis of broad sensory areas showed no difference in the number of activity-labeled neurons across tactile, auditory, or gustatory regions between intact and SW brains. In contrast, a significant enrichment after injury was found in the olfactory network (Fig. 1c). Examination of individual olfactory regions revealed an increase in TRAPed cells across multiple areas (Fig. 1d), including the piriform cortex (PIR) (Fig. 1d,e). Due to high basal activity in the olfactory bulb (OB), we evaluated bulbar activity in independent cohorts of mice treated with a lower dose of 4-OHT (15 mg/kg). Also, OB activity was quantified as the density of TRAPed cells rather than normalized to the total number of labeled cells, as the OB constituted a substantial proportion of the total cell population (∼19.46%), which could obscure region-specific increases or decreases in activity. Relative to intact controls, SW mice exhibited a 2.6-fold increase in bulbar TRAP-labelled neuron density (Fig. 1e), with prominent increases in the granule cell (GCL) and glomerular (GL) layers (Fig. 1e, right), whereas mitral cells showed virtually no labelling (Extended Data Fig. 1a, b). Complementary three-dimensional analysis of cleared brains imaged by light-sheet microscopy further corroborated the elevated TRAP cell density in the OB and in the PIR (Extended Data Fig. 2a-c; Supplementary Movie 1). OB and PIR are two key structures in the brain’s olfactory system. Sensory inputs from the nasal epithelium are first processed in the OB before being relayed downstream to the PIR. These findings identify the olfactory system as a major site of remote activation following visual cortex injury and suggest that specific sensory circuits distant from the lesion may undergo adaptive reorganization.

**Figure 1.**
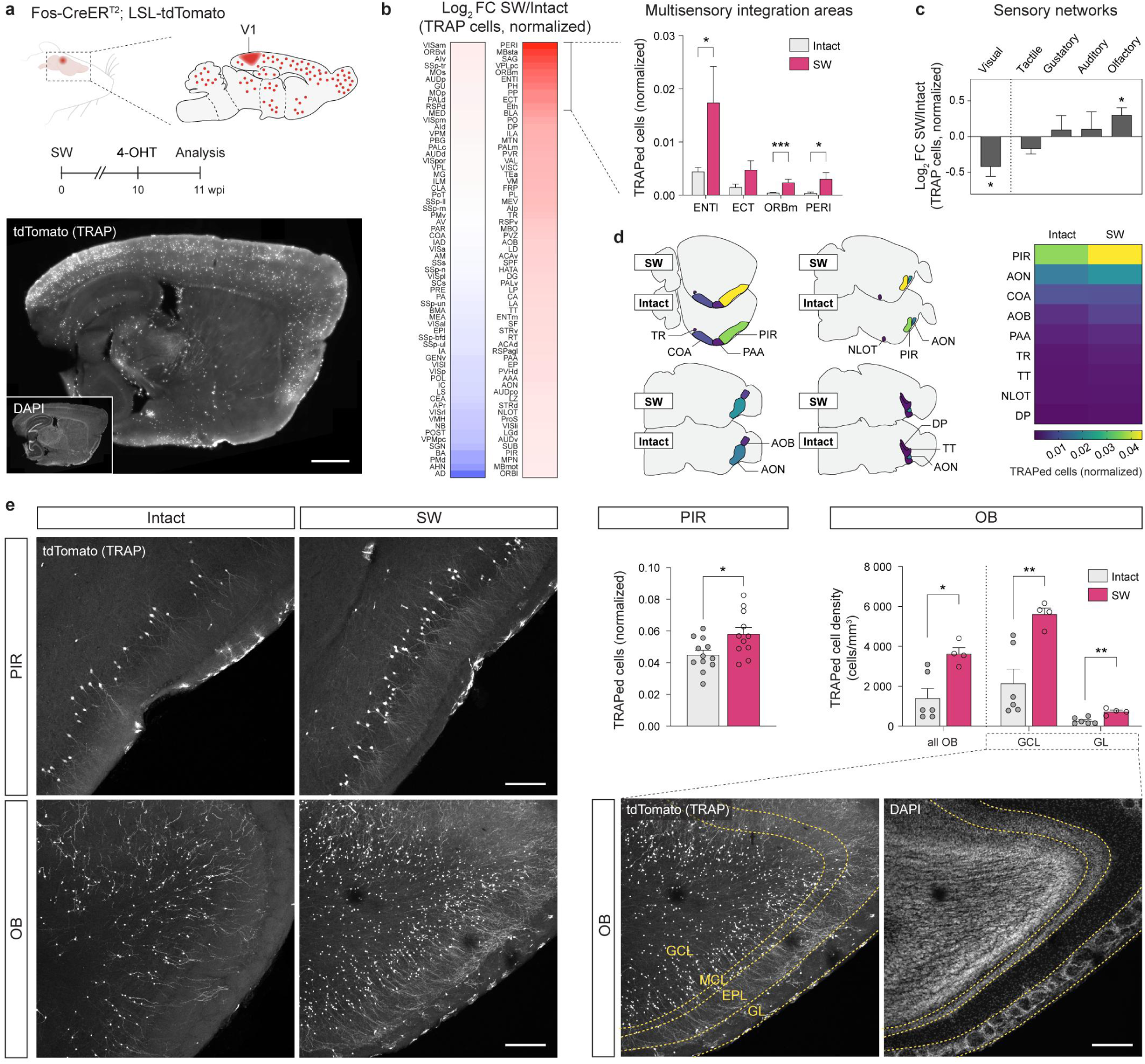
Visual cortex injury increases neuronal activity in olfactory brain areas. **a**, Experimental design. Fos-CreER^T^^2^;LSL-tdTomato mice were subjected to cortical SW in V1 and injected with 4-OHT 10 weeks post-injury (wpi) enabling activity-depending labeling. Representative sagittal section shows TRAPed cells and counterstained DAPI nuclei. **b**, Left, Heatmap displays the log_2_ fold change (log_2_ FC) in normalized TRAP cell counts across brain regions (SW relative to intact controls), plotted on a log_2_ scale to represent proportional increases and decreases with equal visual weighting across orders of magnitude. Right, Normalized TRAP cell counts in multisensory integration areas ranked among the top 10 regions. **c**, Log_2_ FC in normalized TRAP cell counts (SW/Intact) across visual, tactile, gustatory, auditory and olfactory networks. **d**, Schematic sagittal brain sections show olfactory regions color-coded by normalized TRAP cell counts in intact and SW mice. **e**, Representative confocal images and TRAP quantification in PIR and OB of intact and SW mice. OB analysis was performed in mice injected with 15 mg/kg 4-OHT. Bottom right images show delimitation of the OB cell layers based on DAPI signal. Data presented as mean ± S.E.M. and statistical significance was evaluated using Mann-Whitney *U*-tests. \**P* < 0.05, \*\**P* < 0.01, \*\*\**P* < 0.001, \*\*\*\**P* < 0.0001. Sample sizes, *n* = 8 hemispheres from 6 animals (intact) and *n* = 7 hemispheres from 5 animals (SW) (**b-d**); PIR: *n* = 12 hemispheres from 6 animals per group; OB: *n* = 6 hemispheres from 3 animals (intact) and *n* = 4 hemispheres from 2 animals (SW) (**e**). Anatomical abbreviations are listed in Supplementary Table 1. Scale bars, 1 mm (**a**) and 200 μm (**e**). Icons in **a** and **d** created in BioRender and BrainGlobe, respectively.

### V1 injury recruits adult neurogenesis and remodels olfactory bulb connectivity

The OB is continuously supplied with new neurons generated from adult neural stem cells (NSCs) in the subventricular zone (SVZ)^14–16^. Adult-born OB neurons integrate predominantly as granule cells (GCs; ∼90–95%) and to a lesser extent as periglomerular cells (PGC; ∼5– 10%), matching the specific layers where we detected increased TRAP-labelled cell density. Both GCs and PGCs are inhibitory interneurons that modulate the activity of principal projection neurons, mitral cells and tufted cells, thereby shaping olfactory bulb output and odor processing^16^. Adult neurogenesis is thus a potential route through which a focal cortical injury could influence this particular sensory circuit. We therefore examined whether V1 injury alters SVZ neurogenesis using bromodeoxyuridine (BrdU) incorporation assay (Fig. 2a). BrdU pulse labeling was performed one week post-injury, an early time point previously associated with injury-induced activation of adult neurogenesis^17,18^, and revealed increased cell proliferation in the SVZ relative to controls (Fig. 2a-c). This increase was no longer present eight weeks after injury (Fig. 2a, c), revealing the transience of the proliferative response. Cell type-specific marker analysis revealed increased proliferation of both neural stem cells (NSCs) and transit-amplifying progenitors (TAPs) one week after injury, whereas neuroblast volume remained unchanged at this early time point (Extended Data Fig. 3a,b). Consistent with transient NSC activation, NSC density in the SVZ was reduced at one week after injury but was indistinguishable from controls by two months post-injury (Extended Data Fig. 3c). Together, these findings indicate that injury transiently activates the SVZ neurogenic niche without producing a sustained reduction in the NSC pool.

**Figure 2.**
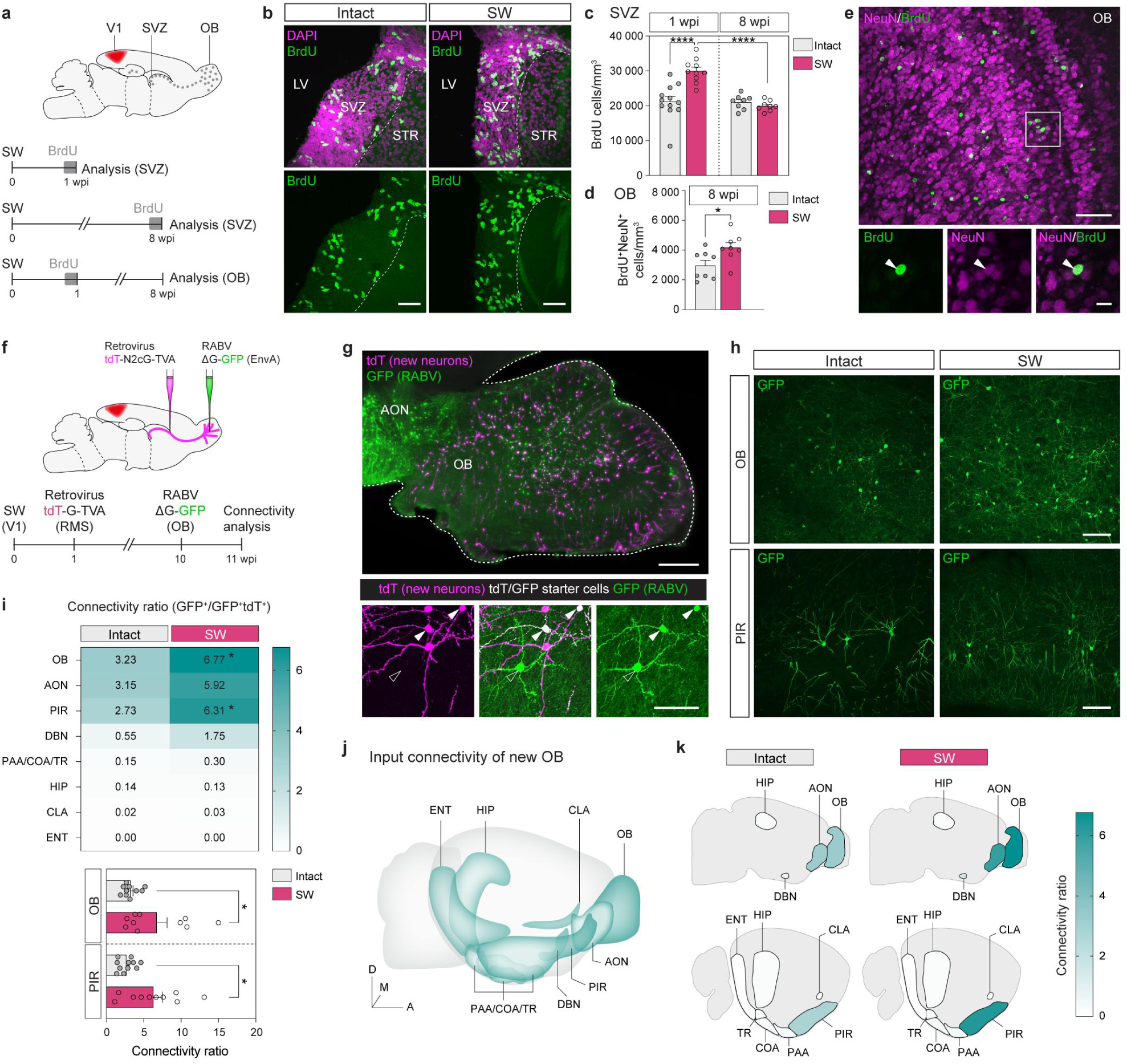
Visual cortex injury promotes SVZ-OB neurogenesis and modifications in adult-born neuron connectivity. **a**, Experimental timelines of BrdU pulse and pulse-chase experiments. **b**, Representative images of the SVZ from intact and SW mice at 1 wpi immunostained for BrdU (green) and DAPI (magenta). **c**, BrdU⁺ cell density (cells/mm^3^) in the SVZ at 1 and 8 wpi. **d**, BrdU⁺NeuN⁺ cell density (cells/ mm^3^) in the OB at 8 wpi. **e**, Representative immunofluorescence image of the OB following a pulse-chase, immunostained for NeuN (magenta) and BrdU (green). Insets show an example of a BrdU⁺NeuN⁺ adult-born neuron (arrowhead). **f**, Monosynaptic retrograde rabies tracing strategy. Injection of a helper retrovirus tdT-G-TVA in the RMS is followed by injection of an EnvA-pseudotyped RABV-dG-GFP in the OB, allows specific targeting of adult-born neurons and monosynaptic restriction of the RABV. **g**, Representative sagittal section shows adult-born OB neurons (tdTomato⁺; magenta), double-labeled starter cells (see inset; tdTomato^+^GFP^+^; white, arrowheads) and their RABV-labeled presynaptic inputs (GFP⁺-only; green, empty arrowheads). **h**, Representative confocal images show GFP⁺ presynaptic neurons in the OB and PIR of mice with a similar number of starter cells. **i**, Heatmap showing the connectivity ratio (number of GFP^+^ cells / number of starter cells) across all identified presynaptic input regions. Bottom, bar/scatter plots indicate connectivity ratio for OB and PIR inputs. **j**, Three-dimensional representation of the input connectome of adult-born OB neurons. **k**, Schematic sagittal brain sections show the afferent regions and respective color-coded connectivity ratio in SW-injured and intact brains. Data presented as mean ± S.E.M. and statistical significance was evaluated using linear mixed effects model (LMM) followed by post-hoc Šídák’s multiple comparisons test (**c**) and Mann-Whitney *U*-tests (**d, i, k**). \**P* < 0.05, \*\**P* < 0.01, \*\*\**P* < 0.001, \*\*\*\**P* < 0.0001. Sample sizes, *n* = 12 hemispheres from 6 animals (intact) and *n* = 10 hemispheres from 5 animals (SW) for 1 wpi; *n* = 8 hemispheres from 4 animals per group for 8 wpi (**c**); *n* = 8 hemispheres from 4 animals in each groups (**d**); *n* = 12 hemispheres from 8 animals (intact) and *n* = 10 hemispheres from 5 animals (SW) (**i-k**). Anatomical abbreviations are listed in Supplementary Table 1. Scale bars, 50 μm (**b** and **e**), 10 μm (**e**, inset), 500 μm (**g**), 50 µm (**g**, inset) and 100 µm (**h**). Icons in **j** and **k** generated using siibra-explorer and BrainGlobe, respectively.

A BrdU pulse-chase experiment showed that the injury-induced increase in SVZ proliferation was followed by a substantial increase in newly generated neurons reaching the OB and expressing the mature neuronal marker NeuN (Fig. 2d,e; Extended Data Fig. 3d). However, the fraction of BrdU-labeled cells that were NeuN-positive was unchanged between injured and control mice (Extended Data Fig. 3e, left), indicating that, among cells generated during the BrdU-labeling period, a similar proportion had acquired a mature neuronal identity at the time of analysis. The increase in newborn neurons occurred without a detectable change in total OB volume (Extended Data Fig. 3e, right), suggesting that this cellular increase did not measurably alter gross OB anatomy. At the molecular level, western blotting of whole-OB lysates revealed increased gephyrin abundance following injury (Extended Data Fig. 3f), consistent with injury-associated remodeling of inhibitory postsynaptic compartments in the OB.

Because the surplus of newly generated neurons might compete for limited synaptic connections, we used a monosynaptic rabies virus tracing strategy targeted to newborn OB neurons^19–21^ to determine whether they establish synaptic connections within the olfactory network. Briefly, a helper retrovirus was injected into the rostral migratory stream (RMS), at 1wpi, to label dividing neuronal progeny with tdTomato while expressing the TVA receptor and the rabies glycoprotein (G) (Fig. 2f). Subsequent injection of EnvA-pseudotyped, G-deleted GFP rabies virus in the OB, restricted the initial infection to TVA-expressing newborn neurons. The rabies G protein supplied in primarily infected neurons (also called starter cells, labeled tdT^+^GFP^+^; Fig. 2g) enabled monosynaptic retrograde spread to their directly connected presynaptic partners, which became labeled with GFP. The absence of rabies G in traced neurons prevented further propagation. OB analysis and brain-wide mapping revealed that the overall distribution of inputs throughout the brain were comparable in intact and injured brains (Extended Data Fig. 4a). Consistent with previous reports, we detected abundant connections within the OB and from the anterior olfactory nucleus (AON), PIR and diagonal band nucleus (DBN)^21,22^. Additionally, we identified sparse connections from three other olfactory regions – cortical amygdalar area, piriform-amygdalar area and postpiriform transition area (COA, PAA, TR) –, the hippocampus (HIP), claustrum (CLA), and entorhinal cortex (ENT) (Fig. 2h,i). All these connections, except from TR, are reported in the Allen Mouse Brain Connectivity Atlas^23^ (Allen Institute for Brain Science, connectivity.brain-map.org; e.g. Experiment 114249084 (COA), Experiment 122641078 (PAA), Experiment 127397469 (ENT), Experiment 286610923 (HIP) and Experiment 187268452 (CLA)). The connectivity ratio for each region was calculated by normalizing the number of GFP⁺ presynaptic input neurons to the number of starter cells in the OB. Quantitative analysis revealed increased presynaptic input from both local OB neurons and the PIR, the principal cortical target of OB output, which also provides feedback projections to the bulb^24^ (Fig. 2h-k). Within the OB, mitral cells – the principal glutamatergic output neurons of the bulb^25^ – represent one potential source of local input to newly integrated interneurons. However, monosynaptic rabies tracing showed that the number of mitral cell inputs per starter cell did not differ between injured and intact mice (Extended Data Fig. 4b). Thus, the increased local OB connectivity observed after injury is unlikely to reflect increased mitral cell innervation and may instead arise from other local presynaptic populations. Consistent with enhanced synaptic drive, BrdU⁺NeuN⁺ neurons in injured mice exhibited higher nuclear c-Fos fluorescence intensity than those in intact mice (Extended Data Fig. 4c). Together, these findings indicate that visual cortex injury increases the production of SVZ-derived neurons and is associated with altered afferent connectivity and functional recruitment of these neurons into OB circuits.

### Injury-induced neurogenesis is associated with refined OB network dynamics

Adult-born GCs regulate olfactory bulb output through reciprocal dendrodendritic synapses with mitral and tufted cells, providing recurrent and lateral inhibition that constrains the spread of excitation and refines odor representations^26^. In addition, cortical feedback amplifies the odor-evoked inhibition of mitral and tufted cells via GC cells^24^. Thus, the increased in both incorporation of adult-born interneurons and piriform afferent connectivity observed after injury may jointly strengthen inhibitory circuit control, promoting sparse, coordinated principal cell activity, and improving odor coding. To examine the functional consequences of these circuit changes, we performed high-density multielectrode array recordings (HD-MEA) in acute OB slices from intact and V1-injured mice (Fig. 3a). Spiking multiunit activity was detected predominantly on electrodes located in the GL, external plexiform layer (EPL), MCL, and at the edge of the GCL (Fig. 3b). Analysis revealed a marked reduction in overall network activity in slices from injured mice, primarily driven by a significantly lower number of active cells on the electrodes (Fig. 3c). In addition, we assessed functional connectivity by quantifying pairwise spike-time correlation coefficients, which measure the synchronicity of neuronal firing between electrode pairs, and node degree (the number of active channels functionally connected to each channel. Despite the reduction in overall activity, pairwise correlation coefficients were increased in slices from injured mice, whereas network node degree was reduced (Fig. 3d). Thus, V1 injury was associated with a smaller and less extensively connected population of active units, but with more strongly correlated activity among the units that remained active.

**Figure 3.**
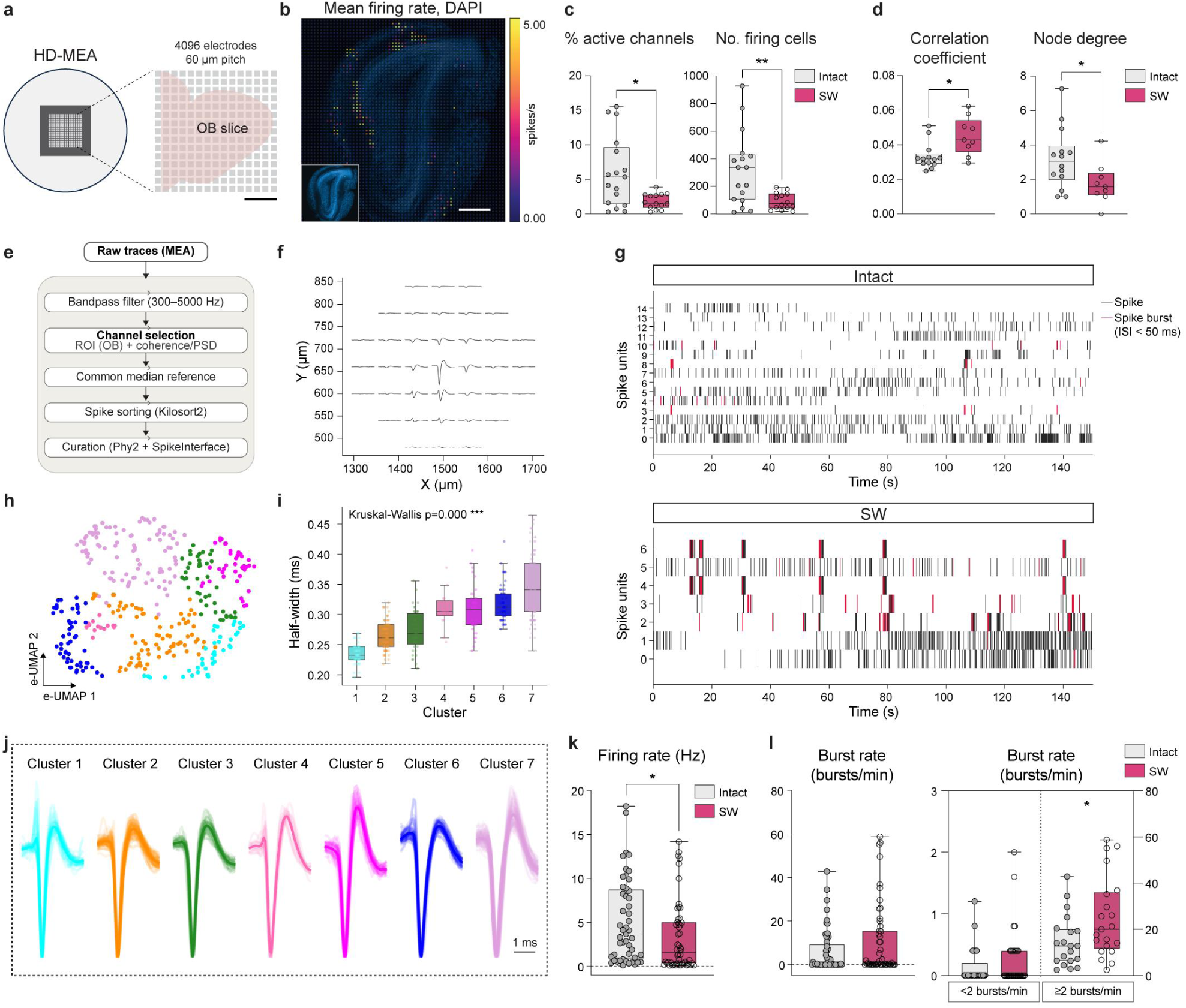
Visual injury is associated with refined OB network dynamics. **a**, Schematic of the high-density multielectrode array (HD-MEA) recording setup. Acute OB slices were prepared from intact controls and SW mice at 12 wpi and placed on a 4,096-electrode array with a 60 µm inter-electrode pitch to record spontaneous extracellular activity. **b**, Representative OB slice with post-hoc DAPI staining, overlaid on its average activity map (mean firing rate heatmap, spikes/s). **c-d**, Electrophysiological parameters assessed include the percentage of active electrodes, number of firing units, pairwise spike-time correlation coefficient, and connectivity node degree. **e**, Spike-sorting pipeline: Raw MEA traces were bandpass-filtered, subjected to region of interest (ROI) selection around OB tissue, and spectral analysis (coherence/PSD) for noise-channel exclusion, common-median referenced, spike-sorted using Kilosort2, and further curated with Phy and an automated SpikeInterface quality-metric filter. **f**, Spatial waveform distribution of a single sorted unit. **g**, Representative raster plots of spike trains from sorted single units, shown for a representative OB slice from intact and SW groups. **h**, Electrophysiological UMAP (e-UMAP) of all sorted single units from intact and SW groups, resolving seven distinct clusters, and (i) the corresponding half-widths (ms) of spike waveforms across clusters 1 to 7. **j**, Representative mean waveforms for each cluster; **k**, Firing rate (Hz) for cluster 7 units in Intact and SW animals. **l**, Burst rate (bursts/min) for cluster 7 in intact and SW animals, shown for units producing fewer than 2 bursts/min (left) and 2 or more bursts per min (right). Data are mean ± S.E.M. Statistical significance across clusters (**i**) was assessed using Kruskal-Wallis tests, Benjamini-Hochberg FDR-corrected, otherwise using two-sided Mann-Whitney *U*-tests (**c-d, k-l**). \**P* < 0.05, \*\**P* < 0.01, \*\*\**P* < 0.001, \*\*\*\**P* < 0.0001. Sample sizes, *n* = 17 slices (4 intact animals) and 14 slices (3 SW animals) (**c-d**) or *n* = 228 cells (13 slices, 4 intact animals), 146 (14 slices, 3 SW animals) units (h–j), or *n* = 49 intact, 52 cluster-7 units (k-l). Scale bars, 1 mm (a), 500 µm (b), 1 ms (j).

To further characterize network activity in the OB, we isolated single units by offline spike sorting (Fig. 3e), enabling extraction of individual neuronal spike waveforms (Fig. 3f). Consistent with the multiunit analysis, single-unit recordings revealed fewer active cells in slices from injured mice, alongside a trend toward increased burstiness (Fig. 3g). We next applied non-linear dimensionality reduction with graph clustering to spike waveforms (*WaveMAP*^27^), identifying seven electrophysiologically distinct waveform clusters (Fig. 3h,j). These clusters differed significantly in waveform half-width (Fig. 3i), peak-to-trough ratio and peak-to-valley duration (Extended Data Fig. 5a). Mapping single-unit locations onto DAPI-stained OB sections revealed preferential spatial distribution across OB layers, with units localized predominantly to the MCL, EPL, GCL, and GL (Extended Data Fig. 5b,c; see Methods). Notably, cluster 7 exhibited the broadest waveforms (Fig. 3i) and putative glutamatergic properties. This cluster was also enriched for units localized to the MCL and EPL (53.13% combined), consistent with a substantial contribution from mitral and tufted cells (Extended Data Fig. 5c). We observed a significant reduction in firing rate in the injured group, potentially arising from differences within cluster 7 (Extended Fig. 5d-e, Fig. 3k). Further characterization of burst rate within cluster 7 did not reveal an overall group difference (Fig. 3l, left). However, analysis of the specific population of cells exhibiting more than 2 bursts per minute revealed a significantly greater number of bursting cells in the injured group (Fig. 3l, right). This data suggest that V1 injury reorganized mitral and tufted cell firing, marked by a sparser, less interconnected OB network in which remaining principal cells fire less often but more synchronously and with greater bursting rates.

To verify the MEA findings specifically in mitral cells, we analyzed the activity of mitral cells by directly expressing the genetically encoded calcium indicator GCaMP6f in these cells and performing two-photon imaging in acute OB slices. We used two complementary labeling strategies: AAV-Syn-FLEX-GCaMP6f delivery to the OB of Tbet-Cre mice^28^ to label mitral and tufted cells selectively (Fig. 4a-c; Supplementary Movie 2) and AAV5-CaMKII-GCaMP6f delivery to the OB of C57BL/6 mice, with imaging restricted to the mitral cell layer (Extended Data Fig. 6a). Both approaches resulted in efficient mitral cell GCaMP6f labeling and yielded comparable results. Consistent with the reduced recruitment detected by MEA recordings, OB slices from injured mice exhibited a smaller fraction of responsive mitral cells compared to intact controls (Fig. 4d; Extended Data Fig. 6b). Responsive cells displayed heterogeneous calcium dynamics that could be classified into three distinct groups: low-activity cells with prolonged transients (<10 events per 5 min; category 1, 42–47%); low-activity cells with brief, sharp transients (<10 events per 5 min; category 2, 19–21%); and highly active cells (>=10 events per 5 min; category 3, 32–39%; Extended Data Fig. 6c,d). The relative representation of these response classes was comparable between groups. Despite the reduced proportion of active cells, mitral cells in the injured mice exhibited a higher frequency of calcium events across both low-(categories 1 and 2) and high-activity response classes (category 3) (Fig. 4f). Moreover, calcium transient amplitude, measured as Δ*F/F₀*, was increased in injured mice (Fig. 4g), showing stronger calcium responses during individual activity events. Thus, V1 injury was associated with sparser recruitment of mitral cells but stronger activity among active cells, consistent with the more coordinated, selectively engaged OB network state observed in multielectrode recordings. Together, these findings suggest enhanced inhibitory control within bulbar circuits, which may increase the contrast between strongly and weakly active principal cell ensembles and thereby support sharpened odor representations.

**Figure 4.**
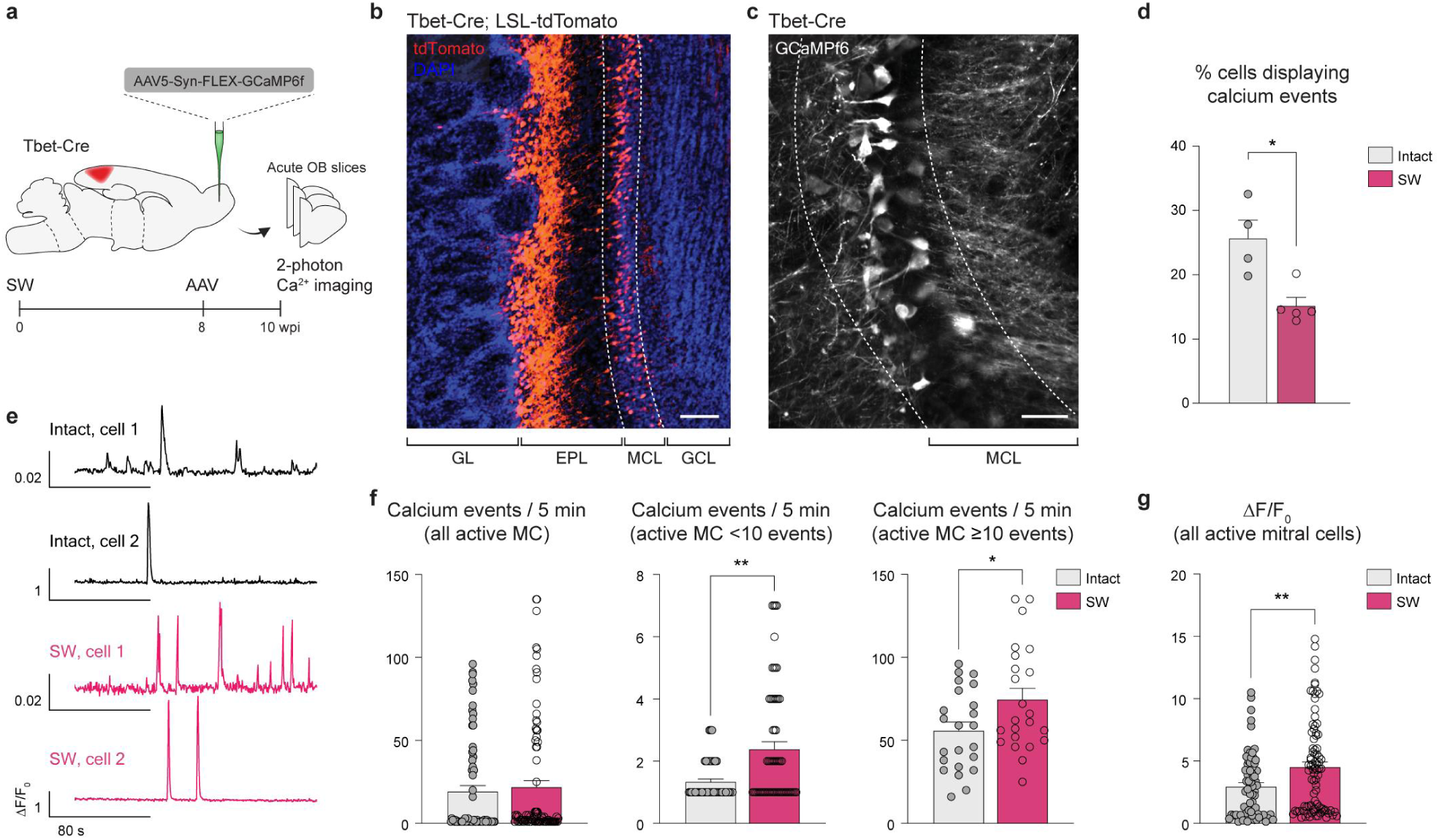
Visual cortex injury potentiates inhibition in OB circuits and sharpens mitral cell calcium responses. **a**, Experimental design for two-photon calcium imaging of mitral cells in acute OB slices. **b**, Representative confocal image of the OB from a Tbet-Cre;LSL-tdTomato mouse shows tdT^+^ mitral cells in the MCL, and tufted cells in the EPL and border with GL, counterstained with DAPI. Dashed lines delimit the MCL. **c**, Representative two-photon microscopy GCaMP6f image of mitral cells in the MCL recorded *ex vivo*. See also Supplementary Movie 2. **d**, Percentage of mitral cells showing calcium events. **e**, Representative traces show relative fluorescence changes (Δ*F/F₀*) over time in mitral cells from intact and SW animals. **f**, Frequency of calcium events (events per 5 min) across all active mitral cells and within low- and high-activity subpopulations. **g**, Mean amplitude of calcium events across all active mitral cells recorded over 5 min. Data presented as mean ± S.E.M. and statistical significance evaluated using Mann-Whitney *U*-tests. \**P* < 0.05, \*\**P* < 0.01, \*\*\**P* < 0.001, \*\*\*\**P* < 0.0001. Sample sizes, *n* = 4 control animals (intact) and *n* = 5 injured animals (SW) (**d**); *n* = 70 cells (14 slices from 4 control animals) and *n* = 85 cells (10 slices from 5 SW animals) (**f**); *n* = 75 cells (14 slices from 4 control animals) and *n* = 89 cells (10 slices from 5 SW animals) (**g**). Scale bars, 100 µm (**b**). 50 µm (**c**).

### V1 injury enhances odor-guided behavior

To establish the behavioral consequences of V1 injury, we first assessed visual performance. In an optokinetic drum test, V1-injured mice showed reduced visual acuity, as indicated by impaired tracking of a rotating black-and-white striped pattern (Extended Data Fig. 7a,b). Consistent with this deficit, V1-injured mice failed to preferentially explore the shallow side over the deep side of a visual cliff apparatus, unlike intact controls (Extended Data Fig. 7a,c; at 10–12 wpi), showing a deficit in depth perception. Thus, a SW injury to V1 induces persistent visual impairment in adult mice.

Next, we examined whether the observed circuit-level changes translated into altered odor-guided behavior. The shift towards sparser, yet more synchronous, mitral cell activity following V1 injury is consistent with an enhanced inhibitory refinement of odor representations in bulbar circuits and improved performance in olfactory tasks^29–32^. In a buried pellet task, V1-injured mice located the hidden food pellet more rapidly than intact controls (Fig. 5a,b), indicating improved performance in an odor-guided search paradigm. We next measured odor detection thresholds using a T-maze assay in which mice chose between a goal arm containing an odor stimulus and an alternative arm containing water. Odor concentrations were progressively increased across consecutive testing days. Compared with intact mice, V1-injured mice detected the odor at lower concentrations (Fig. 5c), demonstrating increased olfactory sensitivity. By contrast, habituation and dishabituation responses to readily detectable odorants were comparable between groups (Fig. 5d). Together, these findings indicate that V1 injury improves olfactory performance under conditions that place high demands on odor detection, enhancing odor-guided search and sensitivity to low-concentration odorants while preserving responses to readily detectable odor cues. Given the persistent visual impairment, these results support the emergence of compensatory enhancement of olfactory function following V1 injury.

**Figure 5.**
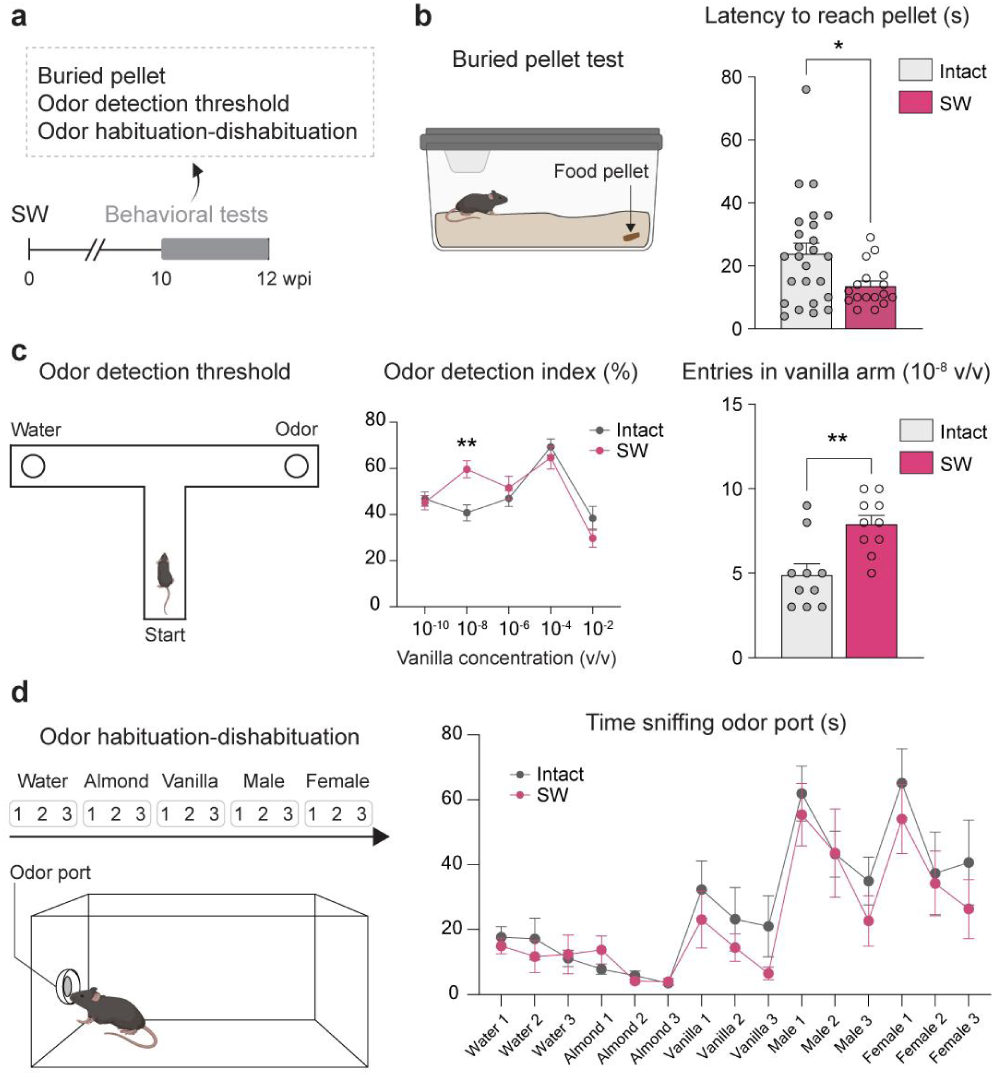
Improved performance in odor-guided behavioral tasks following V1 injury. **a**, Experimental timeline of odor-guided behavioral assays. **b**, Buried pellet test: schematic representation (left) and latency to locate the hidden pellet (right). **c**, Odor detection threshold assay: schematic representation (left), odor detection index across vanilla dilutions (center; 10^−10^ to 10^−2^ v/v) and number of entries into the 10^−8^ v/v vanilla arm (right). **d**, Odor habituation-dishabituation assay: schematic representation (left) and time spent sniffing during each of three consecutive presentations of five odors. Data presented as mean ± S.E.M. and statistical significance was evaluated using Mann-Whitney *U*-tests (**b, c**) and linear mixed models for repeated measured (**d**). Data presented as mean ± S.E.M. \**P* < 0.05, \*\**P* < 0.01, \*\*\**P* < 0.001, \*\*\*\**P* < 0.0001. Sample sizes, *n* = 25 control animals and 17 SW animals (**b**); *n* = 10 animals in each group (**c**); *n* = 10 animals in each group (**d**).

### Adaptive response requires cortical injury and sensory experience

To understand to what extent these olfactory adaptations are driven by visual deprivation versus a general response to cortical injury, we examined mice subjected to an equivalent motor cortex lesion. Unlike V1 injury, motor cortex injury neither altered neuronal activity in olfactory circuits, specifically the PIR (Fig. 6a), nor increased the number of adult-born neurons in the OB (Fig. 6b). Thus, enhanced OB neurogenesis and olfactory function are not general consequences of cortical damage but emerge when cortical injury coincides with an increased reliance on olfactory information.

**Figure 6.**
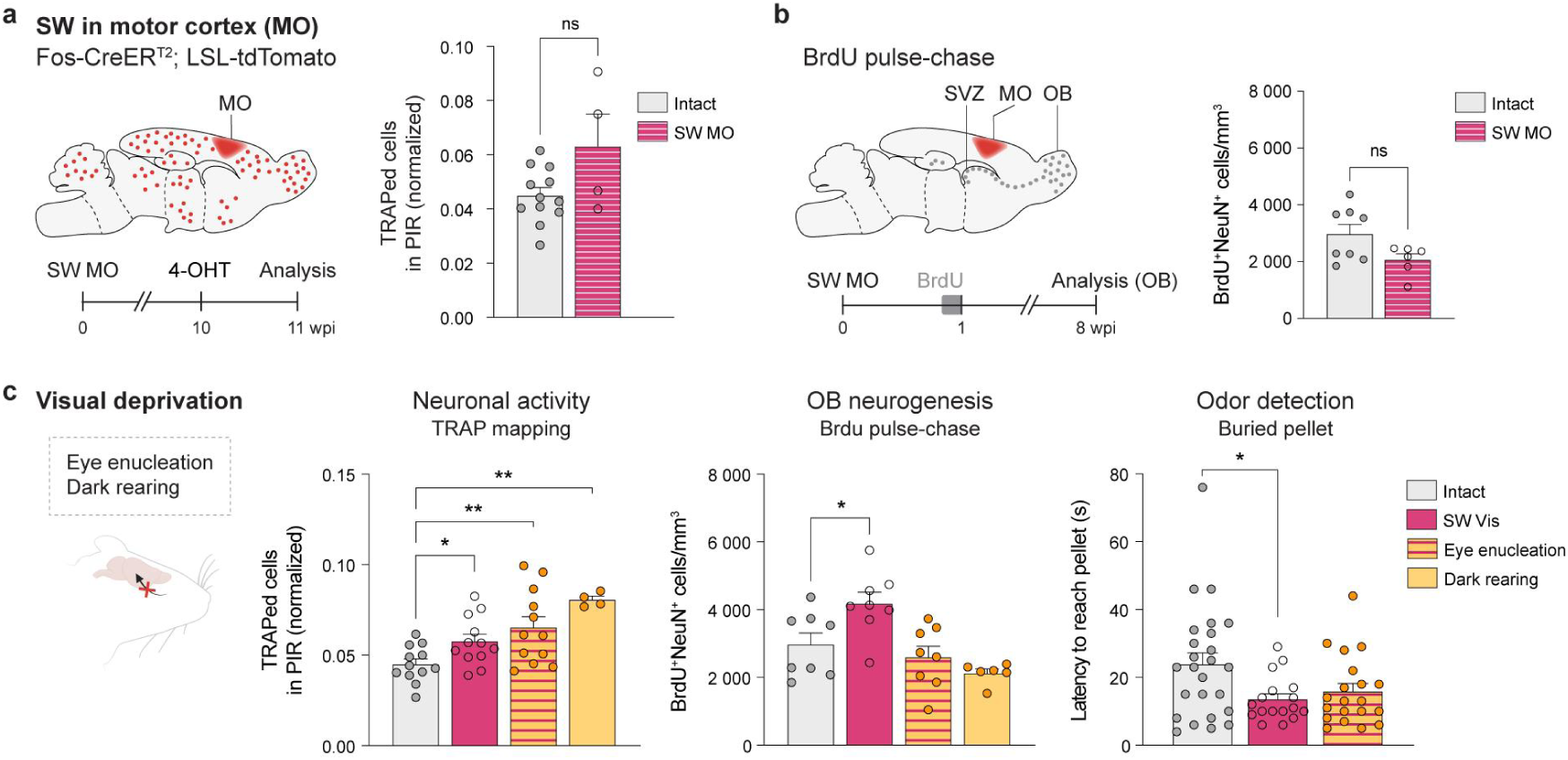
Olfactory neurogenesis and behavioral compensation are not recapitulated by motor cortex injury or visual deprivation. **a**, Left, Experimental design for TRAP mapping. Fos-CreER^T^^2^;LSL-tdTomato mice subjected to motor cortex injury (SW MO) received 4-OHT 10 weeks later to enable activity-depending labeling. Right, Normalized TRAP cell counts in the PIR of intact and SW MO mice. **b**, Left, Experimental design for BrdU pulse-chase experiment. Right, Density of BrdU⁺NeuN⁺ cells (cells/mm^2^) in the OB of intact and SW MO mice. **c**, Comparative analysis of the effects of visual deprivation, induced by eye enucleation or dark rearing, and of visual cortex injury (SW Vis) on PIR activity, adult-born neuron density, and performance in the buried pellet task. Data for SW Vis are reproduced from Fig. 1e, Fig. 2d and Fig. 5b for direct comparison. Data presented as mean ± S.E.M. and statistical significance was evaluated using two-sided Mann-Whitney *U*-tests (**a-b**) or Kruskal-Wallis tests followed by Dunn’s multiple comparisons test (**c**). \**P* < 0.05, \*\**P* < 0.01, \*\*\**P* < 0.001, \*\*\*\**P* < 0.0001. Sample sizes: *n* = 12 hemispheres from 6 mice (intact) and *n* = 4 hemispheres from 4 mice (SW MO) (**a**); *n* = 8 hemispheres from 4 mice (intact) and *n* = 6 hemispheres from 3 mice (SW MO) (**b**); left (PIR activity): *n* = 12 hemispheres from 6 mice for intact, SW Vis and eye-enucleated groups; *n* = 4 hemispheres from 4 mice for the dark-reared group. Centre (adult-born neuron density): *n* = 8 hemispheres from 4 mice for intact, SW Vis and eye-enucleated groups; *n* = 6 hemispheres from 3 mice for the dark-reared group. Right (buried-pellet task): *n* = 25 mice (intact), *n* = 17 mice (SW Vis) and *n* = 20 mice (eye enucleation) (**c**).

In line with this hypothesis, we reasoned that visual impairment may enhance olfactory sampling and sensory reweighting. Consistent with this, V1-injured mice showed increased olfactory engagement during a novel object exploration test as early as one-week post-injury, reflected by a higher sniffing frequency and time spent sniffing, together with prolonged exploration of novel objects (Extended Data Fig. 8a–d). Together, these findings demonstrate that V1 injury is associated with an early increase in olfactory engagement and subsequent enhancement of olfactory circuit function and of odor-guided behavior.

Finally, to isolate the contribution of the olfactory experience from that of the cortical injury, we examined two models of visual deprivation: bilateral eye enucleation and dark rearing (Fig. 6c). Both perturbations increased neuronal activity in olfactory circuits, demonstrating that the mature brain remains capable of engaging remote sensory networks following visual loss, extending earlier observations in younger mice^33,34^. Interestingly, neither model increased OB neurogenesis. However, eye-enucleated mice showed a trend for increased performance in the buried pellet task, but without reaching significance level. Thus, increased olfactory engagement elicited by visual deprivation is insufficient to reproduce the neurogenic and to some extent behavioral adaptations observed after V1 injury. These findings identify cortical injury as a distinct requirement for the observed adaptive response and support a model in which injury-induced neurogenesis, together with olfactory experience, contributes to enhanced odor-guided behavior.

## Discussion

A defining property of complex biological systems is their capacity to compensate for localized failure through systemic reorganization^35,36^. Our findings reveal that the mature brain possesses an underappreciated capacity to reconfigure distant, spared sensory circuits following focal damage, recruiting adult neurogenesis as an active substrate for cross-modal compensation.

Brain-wide TRAP analysis provided an unbiased view of the distributed response to V1 injury. Although the injury was restricted to V1, it was followed by marked reorganization in multisensory integration sites and olfactory regions. Multisensory integration regions are known to participate in blindness-induced cross-modal plasticity^8,37,38^. Such plasticity has been studied most extensively following congenital blindness or early sensory deprivation, in which spared auditory and somatosensory functions can improve through cortical recruitment and refinement of sensory representations^39–41^. Olfactory compensation has received comparatively less attention, although congenitally blind or visually deprived mice exhibit increased sniffing, improved odor-guided behavior, enlarged OBs, and stronger local field potentials in the OB and PIR^33,34,42^. Although brain injury can engage compensatory multisensory mechanisms^43^, whether focal damage to an adult primary sensory cortex actively remodels distant, spared sensory circuits has remained unresolved.

The remote olfactory potentiation was accompanied by increased SVZ–OB neurogenesis. V1 injury induced an early proliferative response in the SVZ involving NSCs and TAPs, in agreement with reports that cortical injury and stroke stimulate SVZ stem cell proliferation and neuroblast production^11,44^ through growth factor, cytokine and chemokine signaling^45^. NSC density was reduced at the early time point but no longer differed between groups at later stages. This recovery may reflect increased NSC quiescence following an initial wave of activation or dedifferentiation of a TAP subpopulation into de novo NSCs^44^.

Injury-induced activation of stem and progenitor cells alone does not explain the sustained increase in adult-born neurons in the OB. Motor cortex injury altered neither olfactory activity nor the abundance of adult-born OB neurons, indicating that cortical injury per se is insufficient to elicit changes in OB. The survival and integration of neuroblasts incoming in the OB are strongly regulated by sensory experience. Odor enrichment and olfactory learning promote the persistence and incorporation of newborn interneurons, whereas odor deprivation has the opposite effect^29,46,47^. Thus, in the absence of enhanced olfactory activity, as observed after motor cortex injury, an injury-induced increase in progenitor production may not translate into sustained incorporation of adult-born OB neurons. By contrast, the increased olfactory sampling observed after V1 injury is consistent with greater reliance on olfactory information, a finding that mirrors well the elevated sniffing reported in congenitally blind mice^42^. Enucleation and dark rearing increased olfactory activity without altering the abundance of adult-born OB neurons, indicating that, in our paradigms, heightened olfactory experience alone cannot expand the integrating neuronal pool in the absence of an upstream injury-induced progenitor response. Together, comparisons with motor cortex injury and visual deprivation indicate that neither cortical injury nor visual impairment alone is sufficient to reproduce the full phenotype. Instead, our findings support a model in which injury-induced stem cell proliferation intersects with experience-dependent olfactory activity to promote the survival and integration of adult-born neurons, enabling adaptive reorganization of a spared sensory circuit. These findings also provide independent support for previous evidence that visual deprivation initiated in adulthood can engage cross-modal plasticity^48^, extending observations otherwise largely limited to congenital blindness or early visual deprivation.

Monosynaptic tracing revealed that adult-born OB neurons generated shortly after V1 injury establish excessive local and corticobulbar connections. Corticobulbar projections onto inhibitory OB interneurons modulate the gain, temporal patterning and correlation structure of mitral and tufted cell activity, thereby contributing to the refinement of odor representations^24,32,49^. This feedback pathway provides state-dependent contextual information that influence early sensory representations in the OB^50^. The coordinated expansion of local and piriform inputs may therefore increase the capacity of adult-born neurons to participate in both local inhibitory processing and context-dependent regulation of OB output. This altered connectivity provides a plausible structural basis for the reorganized OB network dynamics observed by MEA recordings and calcium imaging.

HD-MEA recordings revealed a lower population-level activity, but spike timing more strongly correlated among active electrode pairs. This pattern agrees with the established role of granule cells in synchronizing projection neurons^51^. Mitral cell calcium imaging likewise identified fewer active mitral cells, exhibiting higher event frequencies and larger calcium transients. This response profile may reflect the increased burstiness observed in a putative glutamatergic population identified in HD-MEA. Enhanced inhibitory control onto principal neurons is known to suppress weak or poorly coordinated responses while preserving high-gain responses in selected principal neurons, thereby improving the signal-to-noise properties of olfactory representations and sharpening the separation of relevant odor signals^26,52^. These findings suggest that V1 injury shifts OB activity from a broadly distributed configuration toward a sparser, yet more temporally coordinated, network state. Noteworthy, the increased OB TRAP signal does not conflict with the reduced MEA spiking. TRAP labelling predominantly marked granule cells, whereas TRAP-labeled mitral cells were not detected. This should be interpreted cautiously, as c-Fos may not reliably report mitral cell activity: its induction in mitral cells is stimulus-dependent, and c-Fos-negative mitral cell populations have been reported^53,54^. Conversely, granule cell activity is largely undetected by extracellular arrays. Granule cells exert inhibition through reciprocal dendrodendritic synapses, in which local spine depolarization and Ca²⁺ influx trigger GABA release without generating global somatic action potentials^55,56^. Activity-dependent genetic tagging can therefore capture widespread granule cell recruitment that escapes MEA spike detection.

Our behavioral findings suggest that the structural and functional reorganization has context-dependent consequences rather than producing a uniform enhancement of olfactory function. Injured mice showed increased exploration in the sniffing assay and enhanced performance in tasks requiring detection and localization of low-concentration odor cues. Although greater exploratory engagement may have contributed to performance in these tasks, the lower odor detection threshold also indicates improved access to weak olfactory signals. By contrast, performance in the habituation-dishabituation assay was comparable between injured and intact mice, arguing against a generalized alteration in responsiveness to novel, suprathreshold odors across behavioral contexts. This task dependence is consistent with the variable behavioral consequences reported after experimental manipulation of adult neurogenesis^57,58^. Adult-born interneurons may be particularly influential under conditions that place high demands on sensitivity, pattern separation, learning or the selection of behaviorally relevant signals^30,57^.

Our work reveals the holistic consequences of focal damage to the adult cerebral cortex, revealing significant endogenous plasticity in anatomically distant circuits. Together, our findings establish two principal conceptual advances. First, they demonstrate that cross-modal plasticity follows an acute focal injury to a sensory cortex in adulthood, showing that large-scale sensory reorganization is not restricted to congenital loss or prolonged deprivation. Second, they identify adult neurogenesis as a cellular substrate for sensory compensation, linking the recruitment, survival, and synaptic integration of adult-born interneurons to adaptive circuit remodeling in a spared sensory modality. The present work reveals a previously unrecognized context in which adult neurogenesis is associated with cross-modal compensation and illustrates the capacity of the mature mammalian brain to exploit endogenous cellular resources to generate adaptive network states after injury.

## Methods

### Animals

Wildtype C57BL/6J mice (JAX #000664) were used for histology, electrophysiology, calcium imaging studies and behavioral tests. Fos^CreER^ mice (JAX #021882) were bred with the Ai9 reporter line (RCL-tdT; JAX #007909) for Targeted Recombination in Active Populations (TRAP) in the resulting FosTRAP mice^13^. Tbet-Cre mice (JAX #024507)^59^ were utilized to study mitral cell activity by calcium imaging. Mice were housed and bred in specific pathogen-free conditions at the Institute of Molecular Biotechnology (IMBA) and Institute of Science and Technology Austria (ISTA) in temperature- and humidity-controlled conditions (22 ± 2 °C and 50 ± 10% humidity). Mice were maintained on a 12 h:12 h light:dark cycle with *ad libitum* access to food and water, except stated otherwise. All experiments were conducted in male mice, aged 8–12 weeks at study onset. All animal care, including breeding, husbandry, and experimental procedures, adhered strictly to ethical protocols approved under Austrian and European legislation and the 3R principles.

### Anesthesia and analgesia

All surgeries were performed aseptically under anesthesia with a mixture of fentanyl (0.05 mg/kg; Piramal Critical Care B.V.), midazolam (5 mg/kg; Hameln Pharma GmbH), and medetomidine (0.5 mg/kg; Orion Pharma). After surgery, anesthesia was terminated with atipamezol (2.5 mg/kg; Orion Pharma), flumazenil (0.5 mg/kg; B. Braun Meisungen AG), and buprenorphine (0.1 mg/kg; VetViva Richter GmbH). Meloxicam (1 mg/kg; Metacam, Böhringer Ingelheim Vetmedica GmbH) or Rimadyl (0.15 mg/ml in drinking water; Carprofen, Zoetis Österreich GmbH) was administered as postoperative analgesia for three days.

### Cortical brain injury

SW injury was performed under anesthesia as described in Grade et al. 2022^60^. Following a 2.5-mm-wide craniotomy over the V1, a 1-mm-long and 0.5-mm-deep mediolateral stab wound was made using an ophthalmological lancet (coordinates from lambda: AP: +0.5 mm and ML: ± 2.5 mm). The procedure was repeated in the contralateral hemisphere. SW injury in the motor cortex followed the same procedure with stereotaxic coordinates adjusted to AP +1.25 mm and ML ±1.5 mm relative to bregma. The bone flap was placed back, the skin was stitched and the anesthesia reversed.

### Visual deprivation

Bilateral eye enucleation was performed under anesthesia as previously described^61^. For dark rearing, mice were housed in opaque black polysulfone cages (Tecniplast GM500SUN) for 10 weeks. Dim red light was used for routine animal monitoring, cage changes, and 4-OHT administration to avoid activating the visual system.

### 4-OHT and BrdU administration

30 mg/kg 4-OHT (Sigma) was prepared in 99% ethanol (Millipore) and corn oil (Biozol Diagnostica) at a 1:9 ratio and administered intraperitoneally (i.p.) to FosTRAP mice 8-10 weeks after brain injury (SW Vis or SW MO), eye enucleation or dark rearing. Dark reared mice were returned to standard light conditions 6 days after the 4-OHT administration. In selected experiments (Fig. 1e, as specified in figure legend), 4-OHT was administered at 15 mg/kg, a dose used to facilitate analysis of OB activity given the high Cre-mediated recombination efficiency in this region. All mice were perfused 1 week after 4-OHT injection. For BrdU pulse experiments, 50 mg/kg BrdU (Sigma) was injected i.p. 1 week or 8-10 weeks after visual cortex injury (SW Vis) and perfused 1 h post-BrdU. For BrdU pulse-chase experiments BrdU was i.p. injected 1 week after brain injury (SW Vis or SW MO), eye enucleation or dark rearing. Mice were perfused 8-10 weeks later.

### Monosynaptic rabies virus tracing

To analyze the monosynaptic input onto adult-born neurons generated after injury or visual deprivation, we utilized the rabies virus (RABV) retrograde tracing system, employing the CVS-N2c strain^19,20^. The RABV particles were EnvA-pseudotyped and expressed GFP instead of their own glycoprotein (N2cG). The deletion mutant RABV thus specifically infects cells expressing TVA receptors and its transsynaptic spread is made possible by trans-complementation and N2cG coating in the infected cells. For this, a Moloney murine leukemia virus (MMLV)-based retroviral vector CAG-Tomato-P2A-N2cG-P2A-TVA (1×10^8^ TU/mL) was injected bilaterally in the rostral migratory stream (RMS; 250 nl/hemisphere; from bregma [in mm]: AP +2.25, ML ±0.82, DV −3.5) 1 week after injury or initiation of the visual deprivation. Ten weeks later, the RABV (8.5×10^8^ TU/ml) was injected in both OBs (500 nl/hemisphere; from bregma [mm]: AP 4.35, ML ±0.70, DV −1.5). RABV injection results in infection of tdTomato cells and subsequent propagation of the virus limited to one synapse. Cells sequentially infected by both viruses are identified by GFP and tdTomato double labeling and named “starter cells”, while their first-order pre-synaptic partners receive the RABV particles and thus appear labeled only with GFP fluorescence (Fig. 2g).

In brief, anesthetized mice were head-fixed in the stereotaxic frame (World Precision Instruments, WPI), the skin above the area of interest was cut and the virus was injected at slow rate (1nl/sec) through a drilled burr hole in the skull using an automatic microinjector (Nanoliter 2010, WPI). The skin was stitched back and the anesthesia was reversed.

### Tissue preparation and immunostaining

For immunofluorescence in the brain tissue, mice were deeply anesthetized with ketamin (10 mg/ml Ketamidor, VetViva Richter GmbH) and xylazine (1 mg/ml Xylopan, Vetoquinol Östereeich GmbH) (ket/xyl) and perfused transcardially with phosphate buffered saline (PBS) for 5 min followed by 100 ml of 4% ice-cold paraformaldehyde (PFA, Sigma). Brains were extracted, post-fixed in 4% PFA overnight at 4 °C, then embedded in 4% agarose (Sigma) and sectioned into 70 µm-thick sagittal sections using a vibratome (VT1000S, Leica).

For BrdU labeling, free floating brain slices were subjected to antigen retrieval treatments: citrate buffer (concentration 10X, pH 6.0) (Sigma) for 20 minutes at 96 °C for anti-rat BrdU or 2N HCl (Normaton) for 30 min at room temperature, for anti-rat and anti-mouse BrdU antibodies respectively, before the immunostaining protocol. For immunostaining, brain sections were washed in PBS for 10 min three times and then blocked/permeabilized with 10% normal goat serum and 0.5% Triton X-100 for 2 h at room temperature. Sections were incubated with primary antibodies diluted in blocking and permeabilization solution, for 2 days at 4 °C. The following primary antibodies were used: rat anti-BrdU 1:250 (Abcam), mouse anti-BrdU 1:200 (Sigma), rabbit anti-GFAP 1:500 (Dako Agillent), rat anti-Sox2 1:1000 (Thermo Fisher Scientific), mouse anti-Mash1 1:100 (BD Biosciences), guinea pig anti-Dcx 1:500 (Sigma), rabbit anti-Dcx 1:500 (Abcam), mouse anti-NeuN 1:200 (Sigma), rabbit anti-RFP 1:1000 (Rockland), chicken anti-GFP 1:1000 (Aves Lab), rabbit anti-cFos 1:1000 (Abcam), mouse anti-T-bet (Tbx21) 1:100 (Thermo Fisher Scientific; eBioscience). Next, sections were washed three times in PBS and incubated for 3 h at room temperature with species-specific secondary antibodies: donkey anti-rabbit Alexa Fluor 488 or Alexa Fluor 647 (Thermo Fisher Scientific) or Cy3 (Dianova); goat anti-mouse IgG1 Alexa Fluor 488; goat anti-rat Alexa Fluor 488 or Alexa Fluor 647, donkey anti-rat Alexa Fluor 594; goat anti-guinea pig Alexa Fluor 488 or goat anti-guinea pig Cy3 (Dianova); and goat anti-chicken Alexa Fluor 488 (Thermo Fisher Scientific). Secondary antibodies were diluted 1:500 or 1:1000 depending on the primary antibody concentration. Sections were then counterstained with DAPI (5 μg/ml; Thermo Fisher Scientific) for 10 min, washed three times, and mounted on glass slides with Aqua-Poly/mount (Polysciences).

### Widefield and confocal imaging

For brain-wide analysis of neuronal activity (TRAP) and adult-born neuron connectivity (RABV), one every third section of the brain was stained for GFP, RFP and DAPI and imaged using the Pannoramic 250 FLASH II whole slide scanner (3DHistech) equipped with a CIS VVC FC60FR19CL camera (Vital Vision technology) and a 20x/0.8 plan-Apochromat objective. Images were acquired and visualized using CaseViewer software and Fiji, respectively. For high magnification and high-resolution Z stacks of selected regions, images were acquired using a laser scanning confocal microscope Axio Observer Z1 (ZEISS) equipped with an LSM 800-point laser scanning confocal unit and an Axiocam 705 camera, or with an upright point scanning confocal microscope LSM 800 Axio Imager (ZEISS) equipped with high sensitivity GaAsP detectors and a Hamamatsu ORCA Fusion CMOS camera. A 20x/0.8 Plan Apochromat DIC objective and a 40x/1.3 oil immersion Plan Apochromat DIC objective (ZEISS) were used with z-steps of 2 and 1 µm, respectively. Tiled images were stitched using ZEN Blue software (ZEISS).

### Tissue clearing and light-sheet imaging

FosTRAP mice were transcardially perfused with ice-cold PBS containing 10 U/ml of heparin sodium salt (Carl Roth) followed by 100 ml 4% PFA. After extraction, brains were post-fixed overnight in 4% PFA at 4 °C and cleared following the Fast 3D Clear protocol^62^ with the following modifications: incubations in 50% and 70% tetrahydrofuran (THF with BHT, Sigma-Aldrich) with 0.1% and 0.15% triethylamine respectively (Sigma-Aldrich) (pH 9) were prolonged to 2 h, at 4 °C; incubation in clearing solution was extended to ∼36 h at 37 °C under gentle agitation (100 rpm). The clearing solution consisted of 96% Histodenz, 1.2% Diatrizoic Acid, 2% N-Methyl-D-Glucamine, 0.02% Sodium Azide and 20% Ultrapure Urea dissolved in distilled water (final refractive index [RI] 1.512-1.518). Cleared samples were glued to a 3D-printed sample holder and submerged in an imaging chamber filled with RI-matching Cargille immersion oil (RI 1.512). Images were acquired using a Zeiss Z1 light-sheet microscope with Zeiss Zen software. Samples were imaged in the axial orientation with double-sided illumination using a 5x/0.1 excitation objective and a 2.5x/0.16 detection objective at 1.83 µm/pixel and a z-step size of 4 µm. Neurons labeled with tdTomato were imaged using a 561 nm laser set to 15-20% intensity with 120-125 ms exposure time depending on the sample batch. The autofluorescence channel was acquired using the 488 nm laser line set to 25% power with 150 ms exposure time. Tile overlap was set to 5%. Raw data (.czi) were stitched using Huygens Professional software package (Scientific Volume Imaging) and converted into HDF5 (uncompressed 16-bit unsigned integer) hierarchical format for further processing.

### Western blot

Frozen OBs were defrosted on ice and homogenized with ceramic beads in RadioImmunoPrecipitation Assay (RIPA) buffer supplemented with HALT™ protease inhibitor (100X), following the manufacturer’s instructions (78438, Thermo Fisher Scientific) (2 × 30 s, 4500 rpm). Samples were centrifuged 15,000 × g, 15 min, 4 °C, and supernatants stored at −70 °C. Protein concentration was determined with the Bradford Assay kit (23246, Thermo Fisher Scientific) following the manufacturer’s recommendations: 30 µg protein was mixed with 4x NuPAGE LDS buffer (NP0007, Invitrogen) and 10x NupAGE reducing agent (NP0004, Invitrogen), then resolved by SDS-PAGE and transferred onto PVDF membranes by wet transfer. Membranes were blocked in 5% non-fat dry milk in TBS-T for 1 h at room temperature and incubated overnight at 4 °C with anti-gephyrin antibody (Cell Signaling Technology 14304, 1:1000) or β-actin (MAB1501 Merck, 1:3000) diluted in 5% bovine serum albumin (BSA). After washing three times in TBS-T (10 min each), membranes were incubated with anti-rabbit or anti-mouse HRP-conjugated secondary antibody (Cell signaling 7074 and 7076, respectively), 1:5000) for 1 h at room temperature, followed by additional washes in TBS-T. Immunoreactive bands were visualized using enhanced chemiluminescence (Pierce™ ECL Plus Western Blotting Substrate, 32132X3 Thermo Fisher Scientific) and detected by digital imaging (Biorad Chemidoc). Densitometric analysis of Western blot bands was performed using Fiji (ImageJ, v1.54f)^1^. For each blot, the rectangle tool was used to surround the bands. Band intensity profiles were generated with the Gel analyzing tool in Fiji. The area under each peak was measured after manually drawing a baseline to close each peak, normalized to a loading control of β-actin and the fold change to control groups was calculated.

### Preparation of acute OB slices

Mice were anesthetized with ket/xyl as previously described and transcardially perfused with 20 mL of ice-cold sucrose-based artificial cerebral spinal fluid (suc-ACSF) containing in mM: 210.3 sucrose, 3 KCl, 1.3 MgCl_2_, 2 CaCl_2_, 26 NaHCO_3_, 1.25 NaH_2_PO_4_, and 20 glucose (pH 7.4), bubbled with 95% O_2_-5% CO_2_, and the brains quickly extracted. Horizontal sections (300 μm) comprising the OB were prepared using a vibratome (Leica VT1000S) and were maintained at 32 °C in artificial cerebral spinal fluid (ACSF) containing in mM: 125 NaCl, 26 NaHCO_3_, 3 KCl, 2 CaCl_2_, 1.3 MgCl_2_, 1.25 NaH_2_PO_4_, and 20 glucose (pH 7.4) bubbled with 95% O_2_-5% CO_2_. Slices were used for MEA electrophysiology and calcium imaging in mitral cells.

### Two-photon calcium imaging and analysis

For calcium imaging experiments, C57BL/6J mice received bilateral injections of AAV5-CaMKII-GCaMP6f (Addgene; 4.1 × 10^12^ vg/ml) in the OB. Alternatively, Tbet-cre mice were used in tandem with a Cre-dependent AAV5-Syn-FLEX-GCaMP6f virus (Addgene; 7 × 10^12^ vg/ml) following the same procedure. AAV-CaMKII-GCaMP6f was injected 10-12 wpi, and slice imaging was performed approximately 2-3 weeks later. AAV5-Syn-FLEX-GCaMP6f was injected 8-10 wpi, and imaging was performed 2 weeks later. Acute OB slices were prepared for *ex vivo* calcium imaging in mitral cells.

Two-photon calcium imaging was performed using a SutterInstrument MOM microscope (RAPP OptoElectronic) with a Vidrio Rapid Multi-Region (RMR) scanner, and ScanImage premium software (Vidrio Technologies), equipped with filter sets, and Hamamatsu H11706P-40-A0 uncooled GaAsP PMTs (RAPP OptoElectronic), and Janelia wide path filter design (LP565 R 420-545, T 575-750; BP 525-70; BP 605-70). Fluorescence was collected through a Nikon LWD 16x/0.80 W water-dipping objective (MRP07220, WD 3.0 mm). Excitation was provided by a SPK-ALCOR-920-XSight-2W femtosecond laser (920 nm, <100 fs pulse width, 2 W, 80 MHz) with an acousto-optic modulator for power control and pre-compensation (0 to −60,000 fs²); additional optical components were from Thorlabs. Imaging fields were centered on the mitral cell layer. The mitral cell layer was identified as a distinct row of large triangular-shaped cells (soma area >100 µm^2^) at the outer boundary of the granule cell layer, bordering the external plexiform layer. Imaging was performed in a single optical plane at 2.5 Hz for 5 min per field of view at 32–34 °C under continuous perfusion with oxygenated ACSF. Calcium traces were extracted using the Detect software (GitHub, nsdesai/detect), which integrates image drift correction using the PatchWarp algorithm, and active cell soma and their fluorescence traces identification using constrained non-negative matrix factorization (CNMF). *dF/F_0_* was calculated with *F*_0_ as the lowest value and Fpeak as the highest in each trace, and peaks exceeding *F*_0_ by >10% were included. For cells with fewer than 10 calcium peaks in 5 min, peaks were counted manually; for cells showing >=10 events in 5 min, peaks were detected using a custom Python script.

### Multielectrode array (MEA) recordings and analysis

Extracellular field potentials were recorded using a 2D high-density Complementary Metal-Oxide Semiconductor (CMOS)-based biosensor array (3Brain) comprising 4,096 electrodes with a 60 μm pitch and an active area of ∼3.8 mm^2^. OB slices were prepared, gently positioned on the chip and secured with a platinum harp. Slices were continuously perfused with oxygenated ACSF at 4.5-5 ml/min and maintained at 32-34 °C throughout the recordings. Following an equilibration period of ∼10 min, spontaneous activity was recorded for 2.5 min. Raw data was acquired with a 14 kHz sampling rate per electrode using Brain Wave software (version 5), with a hardware high-pass cutoff at 10 Hz. Raw data were stored in .brw format. BrainWave 5 software was used to define the electrodes overlapping with the tissue (region-of-interest (ROI) tool), to extract the number of active units within. Electrodes with a mean firing rate ≥ 0.5 Hz were defined as active and selected for further analysis. Spike detection and sorting were first performed in Brainwave 5 using the Brain Tissue preset mode, which applies waveform feature extraction and clustering. Functional connectivity was quantified in BrainWave 5 (3Brain AG) via pairwise spike-train cross-correlations across electrode nodes within a 3-ms temporal window, assigning sender, receiver, and broker roles from the resulting directed connectivity patterns.

#### Spike-sorting

Spike sorting was additionally performed using a combination of custom pre-processing pipelines and the unified framework SpikeInterface (version 0.101.2; https://github.com/SpikeInterface/spikeinterface)^63^. Raw extracellular signals were band-pass filtered between 300 and 5,000 Hz using a zero-phase, 4th-order Butterworth filter (forward– backward implementation). Noisy channels were automatically detected and excluded based on inter-channel coherence and channel-specific power spectra. Electrodes were restricted to tissue contacting channels by defining an anatomical region of interest (ROI) using the OB tissue image, which was subsequently co-registered with post-hoc DAPI fluorescence images to map electrodes directly to specific OB layers. The remaining channel signals were re-referenced to the median signal (common median reference). The Kilosort 2 algorithm was then used to detect spikes (threshold of 6 standard deviations) and assign spike-times to individual neurons. Units with an initial firing rate below 0.05 Hz were excluded during sorting. Waveforms and quality control metrics were computed using a channel radius of 200 µm around each unit’s peak channel. All units were manually curated using the Phy GUI (version 1.0.9, https://github.com/cortex-lab/phy). Units that followed the biophysical properties of an action potential generation were included, such as coherent waveform patterns across neighboring channels. Units with high cosine template similarities were merged if considered similar. Positive-spiking units were manually excluded, due to their association with dendrites and axons. Spike units in the final dataset were retained if they exhibited a firing rate > 0.1 Hz, signal-to-noise ratio (SNR) > 2, presence ratio > 0.2, ISI violation rate < 0.2 (Hills contamination estimate, 1.5 ms refractory-period threshold).

#### Burst calculations

Bursts were classified using the inter-spike-intervals (ISIs) calculated between consecutive spikes within each unit’s spike-train. A burst was defined as ≥ 3 consecutive spikes with ISIs below 50ms. Burst rate was expressed as per minute, and units were classified into low- and high-activity subgroups using a threshold of 2 bursts/minute (Fig. 3I)).

#### Spike-unit locations

The spatial positioning of each spike-units waveform was estimated by triangulating its amplitude across the surrounding electrodes (monopolar triangulation within a 200 µm radius). The triangulated position of the max amplitude channel was overlaid onto the region-of-interest image of the OB slice for each recording. The anatomical position of each unit was assigned manually by overlaying coordinates of single isolated units onto the co-registered DAPI image to determine its localization across the OB layers for putative cell type interpretation (Extended Fig. 5b-c).

#### Spike-unit clustering

Spike-units were classified into putative cell-types using WaveMAP^27^. First, the maximum amplitude waveform for each unit was baseline-subtracted and normalized such that its trough amplitude equaled −1. Waveforms were embedded using UMAP (n_neighbors = 20, min_dist=0.1) and partitioned using Louvain community detection. The Louvain resolution parameter was selected by sweeping resolution values from 0 to 3 in steps of 0.2, repeating UMAP + Louvain clustering on random 80% subsamples of the units (25 iterations per resolution) and identifying the elbow of the resulting modularity/cluster-count curve. This selected a resolution of 1.0, which yielded nine putative clusters. Next, clusters whose mean waveforms were highly correlated (r ≥ 0.97; average-linkage hierarchical clustering on a 1 − correlation distance matrix of cluster means) were then merged, yielding a final set of seven clusters.

### Behavioral assessments

All behavioral testing occurred during the light cycle. Mice were single housed for at least one week prior to behavioral testing. For all tests, mice were habituated to the test room for at least 30 minutes before testing. Experimenters were blind to experimental groups.

#### Optokinetic reflex

Visual acuity was assessed using an OptoDrum system (Striatech). Each animal was tested twice. Each session included two positive trials, three negative trials, and a 20-s still-observation period. Acuity thresholds were measured using a staircase algorithm at 99.72% contrast, which adjusted spatial frequency based on the mouse’s optomotor responses and produced stable thresholds around 140-150 cycles. In occasional false-positive tracking events, likely caused by brief nose-poking movements, the trial-threshold parameter was increased to 1.8 to improve scoring accuracy. During testing, mice moved freely on the central platform while the rotating grating was displayed, and visual acuity was defined as the highest spatial frequency that evoked consistent, stimulus-aligned head rotation, determined automatically from the tracked positions of the nose tip and body center.

#### Visual cliff test

Visual depth perception was assessed in a custom-built arena (51.5 x 37 x 44.5 cm; length × width × height) divided into a shallow compartment (28 x 37 cm) and a deep compartment (23.5 x 37 cm). Both compartments were lined with photographic paper displaying a black-and-white checkerboard pattern (3 x 3 cm squares). In the shallow side, the patterned surface was positioned 29.5 cm below the top rim of the arena, whereas in the deep side it was set at a depth of 44.5 cm (producing a 15-cm visual drop). A single continuous sheet of transparent Plexiglas spanned both sides at the level of the shallow floor to provide a uniform walking surface and generate the illusion of a cliff while eliminating tactile drop-off cues^64,65^ The relative orientation of the shallow and deep compartments (left vs. right) was counterbalanced across mice. Each animal was placed individually on the shallow side facing the corner furthest from the experimenter and allowed to explore freely for 5 min under ambient illumination (∼140 lux). Behavior was recorded from an overhead camera and analyzed using TopScan software (Cleversys Inc., Reston, VA, USA, version 3.0) to determine the time spent moving in each compartment.

#### Buried pellet test

The buried pellet test was performed as previously described^66^. Mice were singly housed and food-deprived for 24 h before testing. Each mouse was habituated for 5 min to the test cage (37.5 × 21.5 × 15.5 cm; length × width × height), equipped with a filter top and containing a 3-cm-deep layer of bedding. The mouse was then returned to its home cage while a 1.5 g food pellet was buried beneath the bedding in a corner opposite the starting position. The mouse was subsequently placed in the starting corner, facing the wall, and behavior was video-recorded from the side using a Sony Handycam HDR-CX200 under approximately 120 lux illumination. The trial ended when the mouse located and began consuming the pellet, or after 15 min had elapsed. Latency to locate and reach the pellet was scored. In a subset of trials, latency was scored manually using a stopwatch (Roth).

#### Olfactory habituation-dishabituation

Olfactory habituation-dishabituation task was used to evaluate odor discrimination as follows: (i) odor habituation upon successive presentations of the same odor, (ii) odor dishabituation stemming from increased investigation to a novel odor, and (iii) preference for social olfactory cues^67,68^. Non-social odors were obtained by dipping cotton-tipped applicators in falcon tubes containing water, vanilla (10%), and almond (6%) extracts in distilled water. Social odors were obtained by swiping cotton-tipped applicators in a cage containing group-housed mice. Testing was conducted in a standard mouse cage (GM500 GreenLine Type II L cage, Tecniplast) containing a small opening on one sidewall for easy placement and removal of a cone-shaped stainless steel bottle cap. Cotton-tipped applicators were slid into the sipper tube of the bottle cap before placement into the small opening, to prevent direct access and gnawing of the cotton tip. Odors were presented sequentially starting with water, two non-social odors (vanilla and almond) and ending with two social odors (male then female). Each odor was presented 3 times (2 min each) interspaced by 1-min intervals. Trials were recorded with a GigE camera (acA 1300-600 gm, Basler AG) positioned above the cage. The distance travelled during the session and time spent sniffing the odors were captured using the Ethovision video-tracking system (Ethovision XT 17.5, Noldus, The Netherlands). Active sniffing was scored when the nose was directed at a distance of 1.5 cm or less from the cotton-tip applicator.

#### Odor detection threshold

Odor detection thresholds were evaluated in a T-maze (start arm: 77 cm; choice arms: 31 cm each extending from a 12.5-cm central neutral zone; arm width: 12.5 cm; wall height: 17 cm). Lighting was maintained at 150 lux. Mice were first habituated to the maze by allowing free exploration for 5 min with 4 µl droplets of tap water pipetted onto filter paper discs placed at the distal ends of each choice arm. For testing, vanilla solutions (10^−10^, 10^−8^, 10^−6^, 10^−4^, and 10^−2^ v/v in tap water) were used consecutively. During each trial, one arm contained a 4 µL droplet of the test odor and the other contained a 4 µL droplet of tap water, with odor and water sides alternated between trials. Each mouse was placed in the start arm at trial onset, and behavior was recorded for 5 min using a top-mounted video camera (Panasonic WV-CP294 color CCTV camera, analog composite output, 24 V AC or 12 V DC power supply). Time spent in each arm was scored using a stopwatch (Roth). Choice latency was recorded only once the mouse fully entered a choice arm after exiting the start arm, whether it traversed the neutral zone continuously or paused before making a directional choice. The number of entries to odor arm from the neutral arm was automatically scored using Ethovision video-tracking system.

#### Object investigation and sniffing

Mice were placed in an empty cage without bedding with dim light (LUX = 20) for 5 min for habituation. After the habituation period, a Falcon tube filled with bedding was fixed in the center of the cage and mice were recorded for 5 min. EthoVision XT 18 (Noldus) was used to track the mouse, specifically the mouse nose, and measure object investigation. Parameters measured were: (1) sniffing time: total duration the nose was inside a defined 2-cm circular area around the object, and (2) sniffing frequency: number of nose entries into that area. The explorative sniffing was instead measured manually via slow-motion videos. Sniffing was defined by rhythmic snout/whisker movements while the animal held its head in an elevated or object-directed posture, avoiding moments of crouching or grooming.

Data were analyzed in R software. For the manual quantification, statistical tests were run by modelling the data with a generalized linear mixed model (GLMM) with the lme4 R package. Models were selected based on the smaller AIC and were the following: log-normal for sniffing time; negative binomial for number of sniffing events. Model assumptions were tested using the DHARMa R package and comparisons between conditions at the different time points were tested using the emmeans R package. For the object investigation, conditions at each time point were compared using multiple Mann-Whitney *U* tests. Graphs were generated using GraphPad.

### Image analysis

Analysts were blind to experimental groups.

#### 3D image registration and brain-wide cell quantification

Volumetric images obtained by light-sheet microscopy were resampled by bilinear interpolation from a voxel size of 4 x 1.83 x 1.83 μm^3^ to an isotropic voxel size of 8 x 8 x 8 μm^3^. Resulting images were registered to the Allen mouse brain template^69^ (CCFv3) using the ANTsPy^70^ implementation of the Advanced Normalization Tools (ANTs). ANTsPy’s Symmetric Normalization (SyN) algorithm was applied as it combines both an affine and a deformable registration strategy with the mutual information as optimization metric. Cell detection was performed by applying a Laplacian of Gaussian (LoG) filter^71^ using the implementation in the Python package scikit-image^72^. To match expected soma size, we constrained the standard deviations of the Gaussian kernels to the measured cell dimensions. Parameter tuning identified a local-maxima threshold of 0.075 as optimal for our dataset. After detection, double hits in the axial (z) direction were corrected by merging detections within an anisotropic neighborhood of 96 × 16 × 16 μm (12 × 2 × 2 voxels). Finally, we applied the inverse deformation field to map the brain atlas regions to the acquired images and extract the number of cells in each region. The source code of the image registration pipeline is publicly available in https://github.com/BrainImageAnalysis/Mouse_Brain_Cell_Count.

#### 2D image registration and brain-wide cell quantification

To map tdTomato-labeled neurons (TRAP) throughout the brain in images obtained from brain sections, we followed the QUINT workflow^73^. Sections were sampled systematically, collecting every fourth slice throughout the mediolateral axis of each hemisphere. Whole-section images were acquired in two channels for DAPI and tdTomato. QuickNII and VisuAlign tools were used to align the imaged brain slices to the Allen Mouse Brain Atlas Common Coordinate Framework version 3 (CCFv3; 2017). tdTomato-positive cells were segmented using ilastik^74^. The ilastik classifier was trained on manually annotated examples to distinguish neuronal morphologies from non-neuronal morphologies (e.g., meningeal cells and reactive glia in the SW; Extended Data Fig. 9a,b). Objects classified as non-neuronal cells were excluded from downstream analysis. Nutil was used to quantify labeled neurons relative to the reference brain atlas. From the output tables, root, parent, and unassigned atlas regions were excluded, as were the ventricular system, fiber tracts, and areas that could not be reliably aligned or were not consistently represented across mice (pons, medulla, and cerebellum). For each designated brain region, labeled cell counts were then normalized to the total number of labeled cells across all remaining included regions within each brain, thereby accounting for inter-animal variability in Cre recombination efficiency. In addition, TRAP-labeled neurons in the PIR and OB were quantified using an independent, region-of-interest-based approach. OB and PIR ROIs were delineated based on anatomical landmarks in DAPI-stained images. tdTomato-positive cells within each ROI were counted using an automated Fiji macro. Regional activity was expressed as the number of TRAP-labeled cells within each ROI divided by the total number of TRAP-labeled cells in the corresponding section. Additionally, cell density was calculated. This complementary approach was performed using four anatomically equivalent sections per mouse.

#### Neurogenesis analysis

Cell quantification in the SVZ and OB was performed using the Cell Counter plugin in Fiji. Cell density was calculated by dividing the number of positive cells by the total imaged area. The SVZ was delineated on maximum intensity projections of the DAPI channel. Neural stem cells were defined as GFAP^+^Sox2^+^BrdU^+^ triple-positive cells. Neuroblast volume in the SVZ was quantified using the Surface tool in Imaris software (Oxford Instruments). Adult-born neurons in the OB were identified as NeuN^+^BrdU^+^ double-positive cells in BrdU pulse-chase studies. All quantifications were performed on 3-4 equally spaced sagittal slices per brain hemisphere.

#### Connectivity analysis

For brain-wide analysis of rabies-traced connectivity in the OB, every third brain section was collected. Inputs to adult-born neurons in the OB were quantified from slide scanner images. ROIs were manually delineated based on anatomical landmarks in DAPI images. GFP^+^ cells were segmented using ilastik software and counted with a Fiji macro. Double-labeled starter cells (GFP^+^Tomato^+^) were counted in Fiji using the cell counter plugin. GFP-only cells represented first-order connections. The connectivity ratio was calculated by dividing the number of GFP-only cells in each brain region (per hemisphere) by the total number of starter cells in the OB of that hemisphere.

### Statistical analysis

All statistical analyses were performed in GraphPad Prism (GraphPad Software, San Diego, CA, USA) unless otherwise stated. Outliers were identified using the ROUT method in Prism. Statistical tests are defined in respective figure legends.

Data from the investigation/sniffing assay were analyzed in R software. Statistical tests were run by modelling the data with a generalized linear mixed model (GLMM) with the lme4 R package^75^. Models were selected based on the smaller AIC and were the following: log-normal for sniffing time; negative binomial for no. of sniffing events; Gamma for investigation time; negative binomial for investigation events. Model assumptions were tested using the DHARMa R package^76^ and comparisons between conditions at the different time points were tested using the emmeans R package^77^. Graphs were generated using the ggplot2 package^78^.

Data from the odor habituation-dishabituation assay were subjected to linear mixed modelling using the lme4 package in R 4.5.2 (R Core Team, 2025). Primary predictors of interest (fixed effects) were Group (Intact vs. SW), Odor, and Trial. Mouse identity (MouseID) was included as a random effect to account for inter-individual variance. The model was estimated using restricted maximum likelihood estimate (REML), with the significance of fixed effects tested using Type II Wald *F* tests with Kenward-Roger degrees of freedom.

Data are presented as mean ± SEM. Statistical significance was set at *P* < 0.05 (*), *P* < 0.01 (**), *P* < 0.001 (***) and *P* < 0.0001 (****). For statistical reporting, each animal was treated as one biological replicate (n = 1) in behavioral studies; each hemisphere was treated as *n* = 1 in neuronal activity and neurogenesis analyses; each slice or isolated unit (neuron) constituted *n* = 1 in MEA analyses; and each cell (trace) was treated as *n* = 1 in calcium imaging studies, derived from multiple animals as described in respective figure legends.

## Acknowledgements

We thank to the GMI/IMBA/IMP Scientific Service Units, in particular the BioOptics facility (Karin Aumayr and team) and Comparative Medicine (Daniela Pollak and team), for their outstanding support. We are grateful to the VBCF core facilities (Preclinical Phenotyping and Histology) for their support. We thank András Aszodi for the support in the statistical analysis. We thank the Mechanical Engineering Center for designing and manufacturing the visual cliff apparatus. We thank Anna Smolka for technical support, Magdalena Krubner for generating the QUINT analysis scripts, and the Grade laboratory for their help and discussions throughout this project.

## Author contributions

S.G. conceived and administered the project. O.C. and S.G. conceptualized the project and the experimental design. O.C. performed all the experiments except those next stated. S.M. performed the analysis of mitral cell inputs and of investigation/sniffing datasets, M.Z performed the spike sorting analysis and waveform classification, M.M. performed the odor habituation-dishabituation assay and partly the investigation/sniffing tests, S.V.P. performed the tissue clearing and light-sheet imaging, M.D. analyzed the light-sheet imaging data, S.H. performed the two-photon imaging acquisitions, M.N.G.A. performed the western blot studies, L.P. performed the optokinetic drum assay, T.D. performed dark rearing studies together with O.C. and S.M.. S.G. H.S., J.K. and E.T. provided supervision. S.G. and O.C wrote the manuscript with input from all authors.

