## Supplementary Figures for "Adaptive brain rewiring after brain injury via adult neurogenesis"

### Extended Data Figure 1

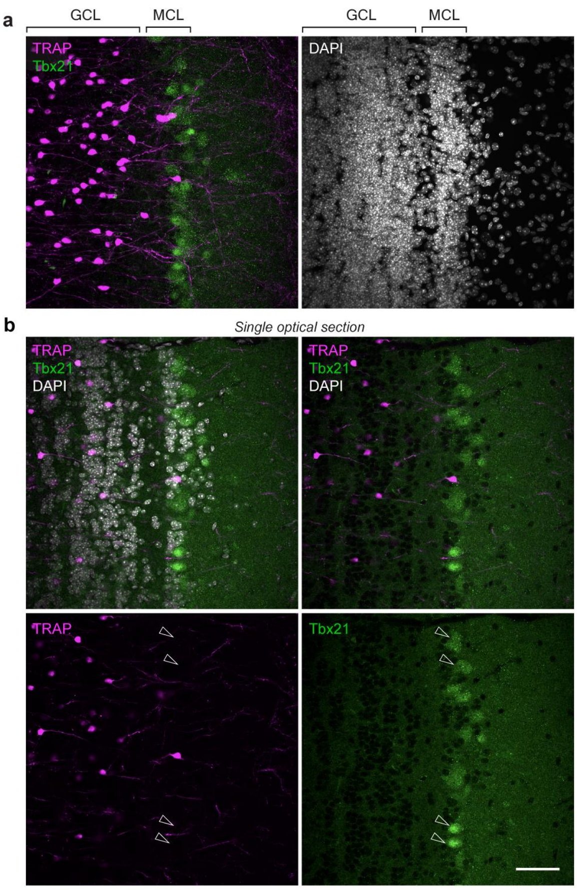

Extended Data Figure 2

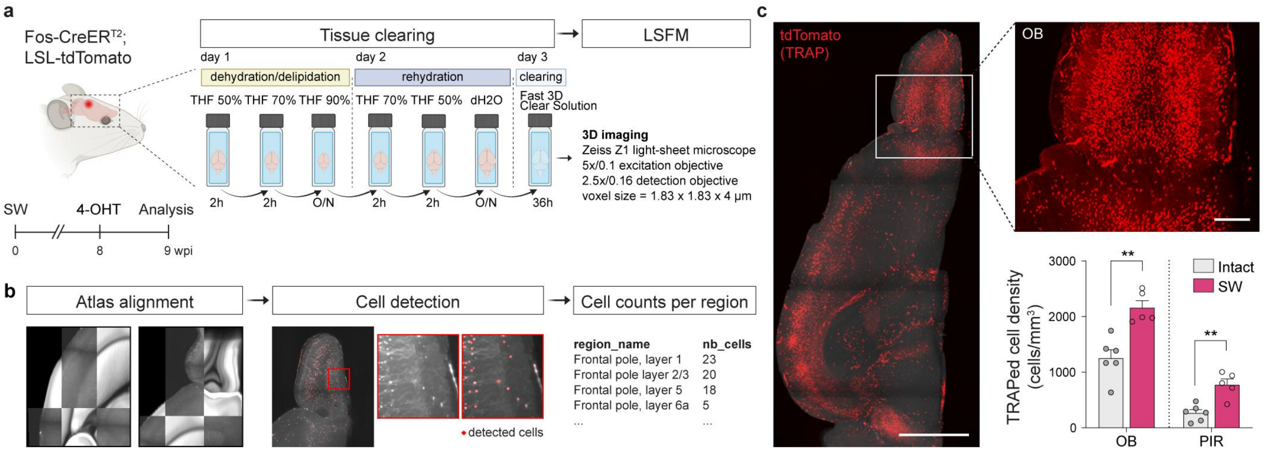

### Extended Data Figure 3

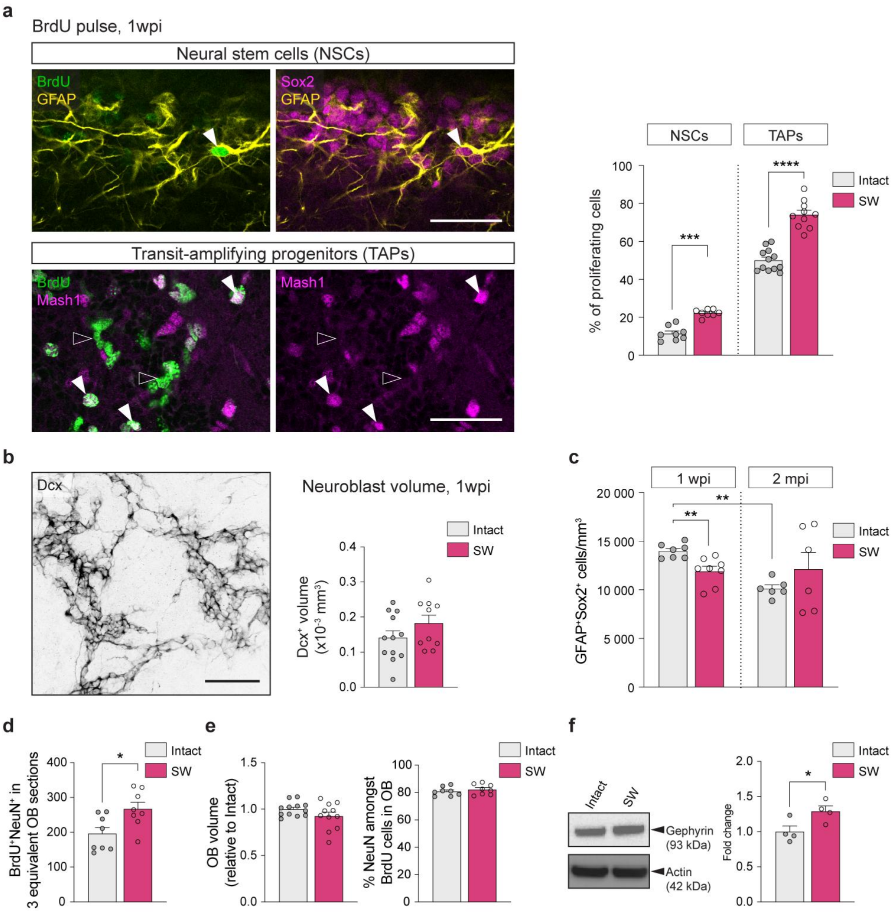

### Extended Data Figure 4

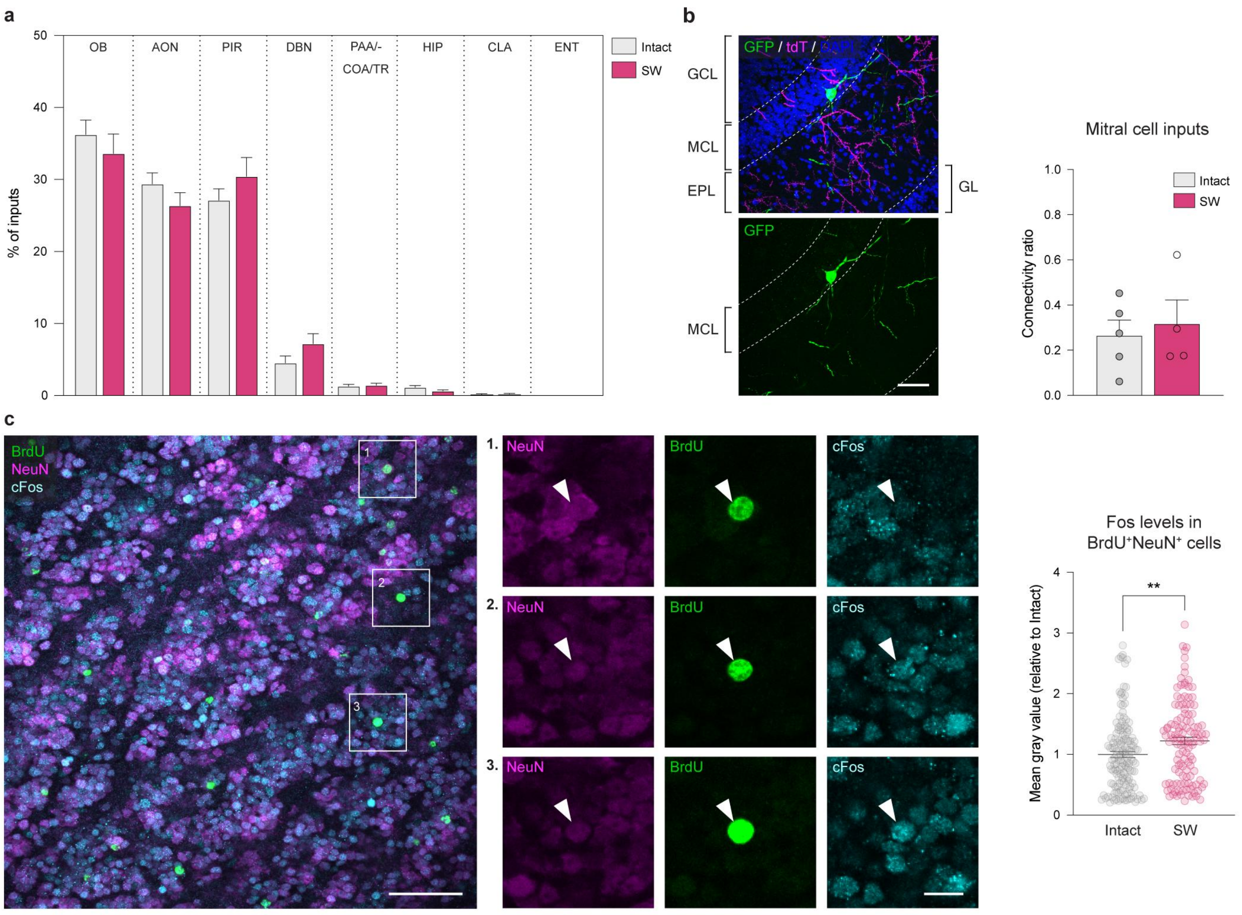

### Extended Data Figure 5

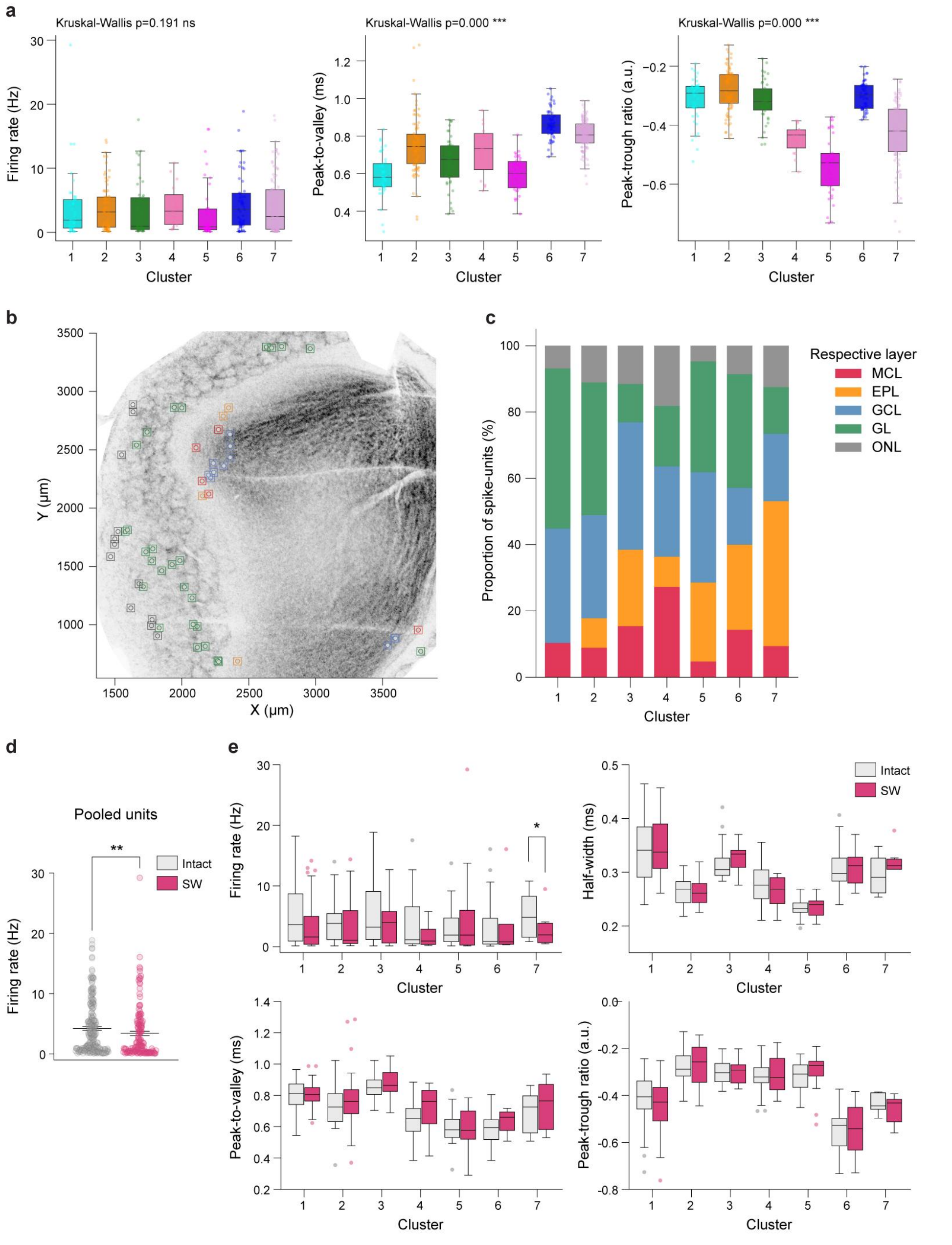

### Extended Data Figure 6

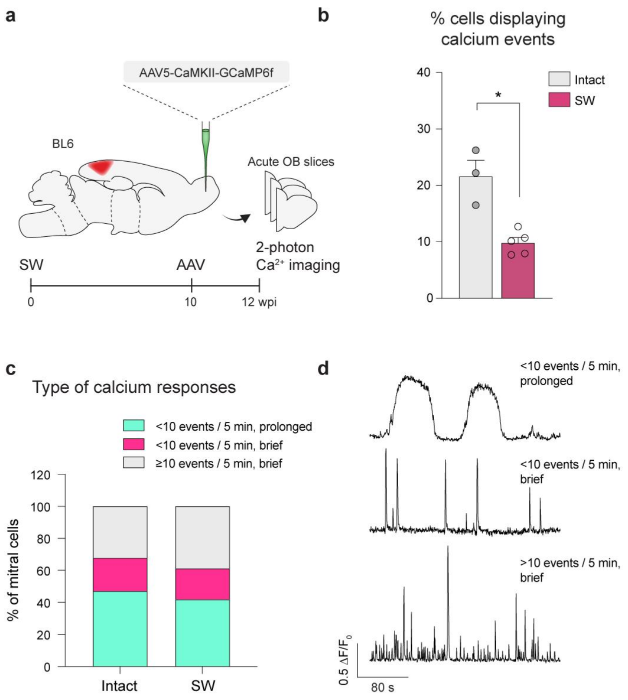

### Extended Data Figure 7

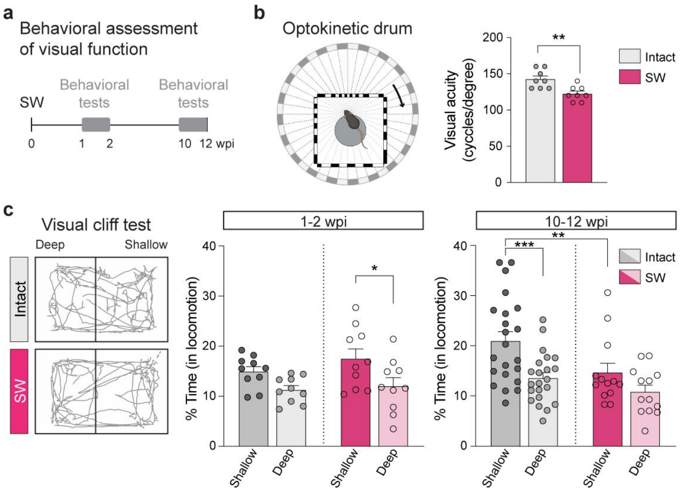

### Extended Data Figure 8

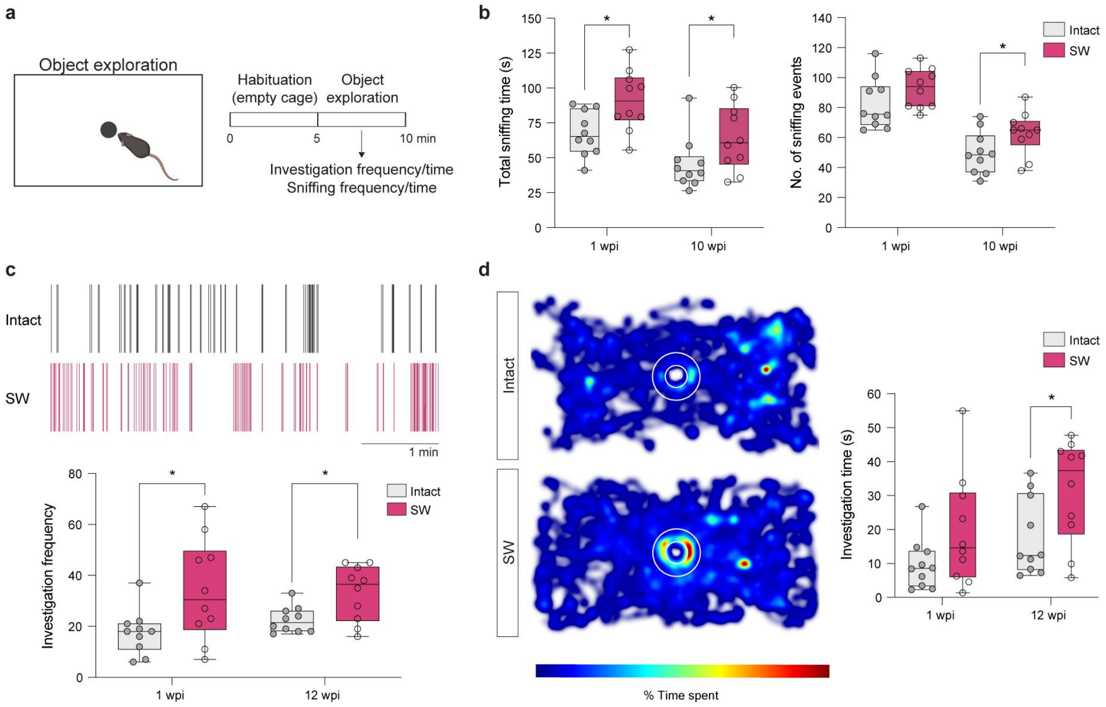

Extended Data Figure 9

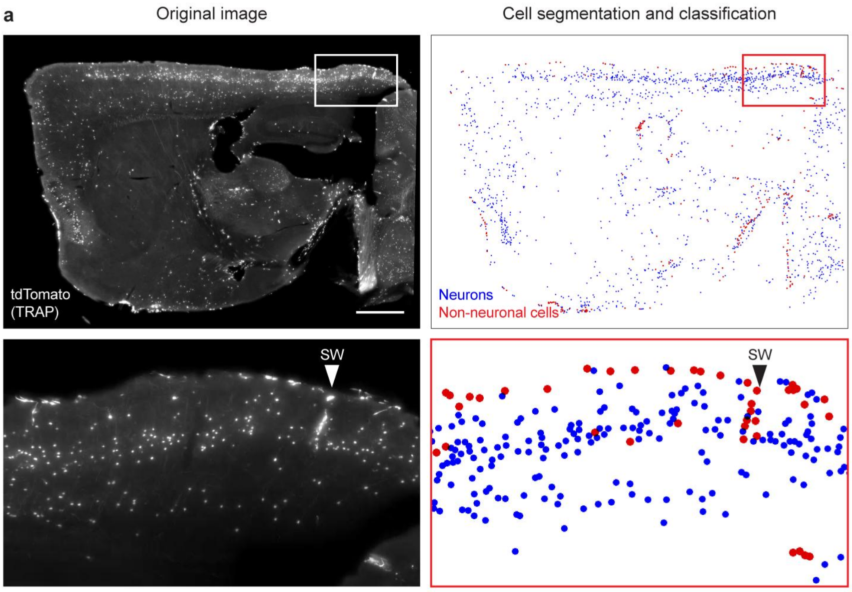
